# A critical analysis of plant science literature reveals ongoing inequities

**DOI:** 10.1101/2022.10.15.512190

**Authors:** Rose A. Marks, Erik J. Amézquita, Sarah Percival, Alejandra Rougon-Cardoso, Claudia Chibici-Revneanu, Shandry M. Tebele, Jill M. Farrant, Daniel H. Chitwood, Robert VanBuren

## Abstract

The field of plant science has grown dramatically in the past two decades, but global disparities and systemic inequalities persist. Here, we analyzed ~300,000 papers published over the past two decades to quantify disparities across nations, genders, and taxonomy in the plant science literature. Our analyses reveal striking geographical biases—affluent nations dominate the publishing landscape and vast areas of the globe having virtually no footprint in the literature. Authors in Northern America are cited nearly twice as many times as authors based in Sub-Saharan Africa and Latin America, despite publishing in journals with similar impact factors. Gender imbalances are similarly stark and show remarkably little improvement over time. Some of the most affluent nations have extremely male biased publication records, despite supposed improvements in gender equality. In addition, we find that most studies focus on economically important crop and model species and a wealth of biodiversity is under-represented in the literature. Taken together, our analyses reveal a problematic system of publication, with persistent imbalances that poorly captures the global wealth of scientific knowledge and biological diversity. We conclude by highlighting disparities that can be addressed immediately and offer suggestions for long-term solutions to improve equity in the plant sciences.

**SIGNIFICANCE STATEMENT:** We analyzed ~300,000 papers published over the past two decades to quantify global, gender, and taxonomic disparities in plant science. Our analyses reveal striking geographical biases that are correlated with national affluence. Gender imbalances were also evident, with far more papers led by authors with masculine names than authors with feminine names. Lastly, we identified substantial taxonomic sampling gaps. The vast majority of surveyed studies focused on major crop and model species and the remaining biodiversity accounted for only a fraction of publications. Taken together, our analyses represent an important addition to the growing conversation about diversifying and decolonizing science.

## INTRODUCTION

Plant science research is accelerating at a rapid pace. New technologies and expanding infrastructure have opened the door for cutting-edge research to be conducted at monumental scales. Despite this noteworthy growth, access to resources is not evenly distributed across the globe and recent studies have revealed striking participation gaps and longstanding disparities tied to colonialism, economic inequality, and systemic biases (1–6). Plant science, which, for the context of this study, we define broadly as *any research investigating an organism that performs photosynthesis*, suffers from acute historical exclusion and ongoing underrepresentation of marginalized identities (7). In Northern America, associations between plant science and agriculture with colonialism, slavery, and the exploitation of migrant workers (8) have contributed to a notable lack of diversity in the discipline. Global economic disparities, established under imperial colonialism and perpetuated through modern eurocentric frameworks, further exacerbate underrepresentation of diverse perspectives in plant science (9, 10, 3). Researchers working in low-income countries and under-resourced institutions face multiple barriers to participating in plant science research, including limited funding opportunities, reduced access to cutting-edge technologies and infrastructure, and exclusion from collaboration networks (5,11). In the field of plant genomics for instance, few projects have been led by researchers in the Global South, despite the striking biodiversity and extensive local botanical knowledge within these regions (3). These dynamics are reinforced by a eurocentric framework that centers English language standards, Latin binomial naming conventions, and reductionist thinking. Coupled with historical and ongoing expropriation of plant germplasm from the Global South, this has resulted in a system that unjustly benefits certain individuals and excludes others. A first step of addressing these inequalities is to quantify patterns of participation in plant science.

Both race and gender compound with global economic disparities to generate emergent barriers for people of color and individuals with marginalized gender identities (2, 12). For example, women of color are uniquely oppressed across multiple axes in ways that amount to more than the sum of their racial and gender identities (12). Although our analyses do not address race directly, we explore global patterns with links to imperial colonialism that cannot be understood without an acknowledgement of race and the persistent oppression faced by people of color, especially Black and Indigenous communities. Our analyses address patriarchy, sexism, and gender dynamics more directly. Patriarchy can be described as a way of living that privileges all men over women and some men over other men, and the politics of patriarchy can be understood as “the politics of domination – a politics that rationalizes inequality” (13). Systems of patriarchy vary in their manifestation and severity across the globe, but are pervasive and have infiltrated all levels of society including scientific research (14–17). While self-identified women are not excluded from the field of biology as a whole, they are often excluded from prestigious tenured and editorial positions as well as collaboration networks (7, 18–20). Studies suggest that gender biases also exist in hiring, publication, and funding decisions (6, 19, 21–25). These inequities impact academic currency on job and funding markets and further exacerbate imbalances in academia. Quantifying the extent and patterns of gender bias in plant science is an important step in creating a more equitable discipline.

Despite noteworthy efforts made towards cataloging all life, research attention has not been equally distributed across study systems and many species remain underexplored. In plant genomics, for example, there are substantial taxonomic gaps–multiple clades lack a reference genome assembly while other clades have dozens of sequenced species (3, 26, 27). These findings suggest that research attention has been disproportionately directed towards a few select species with agricultural and economic relevance to modern society. Focusing on these elite crop and model species has enabled noteworthy scientific breakthroughs and agricultural innovations, but it has come at the cost of exploring the rich biodiversity of wild plants and regionally important crops. With species extinction rates at an all-time high (28, 29), much of this biodiversity could be lost before it is understood scientifically. Participation gaps likely contribute to taxonomic sampling gaps in complex and context dependent ways. For example, the exclusion of Indigenous perspectives from science has removed valuable knowledge of local biodiversity and diverted resources away from regionally important plants (30). Together, these factors exacerbate the patriarchal and eurocentric system of publication, and result in a body of literature that poorly represents the global wealth of biological diversity and knowledge.

To better understand the changing global landscape of plant science research and quantify patterns of underrepresentation, we conducted a large-scale bibliometric analysis of nearly 300,000 papers published across the past two decades of plant science research. Our analyses are framed from the perspective of the first axiom of Ardila-Mantilla which states that scientific potential is “distributed equally among different groups, irrespective of geographic, demographic, and economic boundaries” (31). If we take such a statement to be the null hypothesis, then disparities in educational advancement and promotions, funding, or publication and citation rates indicate that other factors, like oppression, have created historical and contemporary biases in science. To test this hypothesis, we identified the demographic features (e.g., nationality and gender) associated with high publication and citation rates and quantified taxonomic sampling gaps and regional differences in focal organism choice to explore associations between participation gaps and study organisms. We examined how these dynamics change over time and space to identify areas that are improving, stagnant, or regressive. We close by discussing the need to dismantle oppressive systems in the plant sciences, improve equity, and how such changes will ultimately advance the field in the coming decades.

## RESULTS

We compiled a database of 296,447 plant science papers published between 2000 to 2021. Papers were sourced from a representative set of 127 plant science journals based in 26 different nations across 5 continents, covering 21 different subspecialties. We included both society and for-profit journals in our analyses, with open access, hybrid, and subscription publishing models (see Supplementary Dataset S1 for journal information). The database we assembled does not capture the entire breadth of plant science and related fields as many regional and subject specific journals are not included here. As such, these analyses represent an important but non-exhaustive step towards quantifying inequities in plant science.

### Geographic disparities in publication and citation rates

To gain insight into the global landscape of plant science research, we summarized geographic differences in publication and citation numbers. Vast areas of the world have virtually no footprint in the plant science literature over the past two decades (Figure 1A) and publication rate is tightly correlated with national affluence. On a continental level (Supplementary Figure 1A), nearly one third (27%) of all papers were led by authors based in Europe, another 18% were led by authors in Northern America, and 37% by authors in Asia. The remaining 17% of publications were led by authors distributed across Africa, Latin America, and Oceania. Within each continent, authors were further consolidated into distinct hubs of research activity, with the USA, China, and Western Europe dominating the plant science landscape (Figure 1A). National publication rates were highly correlated with Gross Domestic Product (GDP) (R^2^=0.75, F_1,140_=213, p=3.18e-43) (Figure 1B) and investment in research and development (R^2^=0.83, F_2,117_=295, p=2.08e-46) (Figure 1C and Supplementary Figure 2A). Both relationships follow a power law, as reflected by a linear relationship in logarithmic plots.

**Figure 1.**
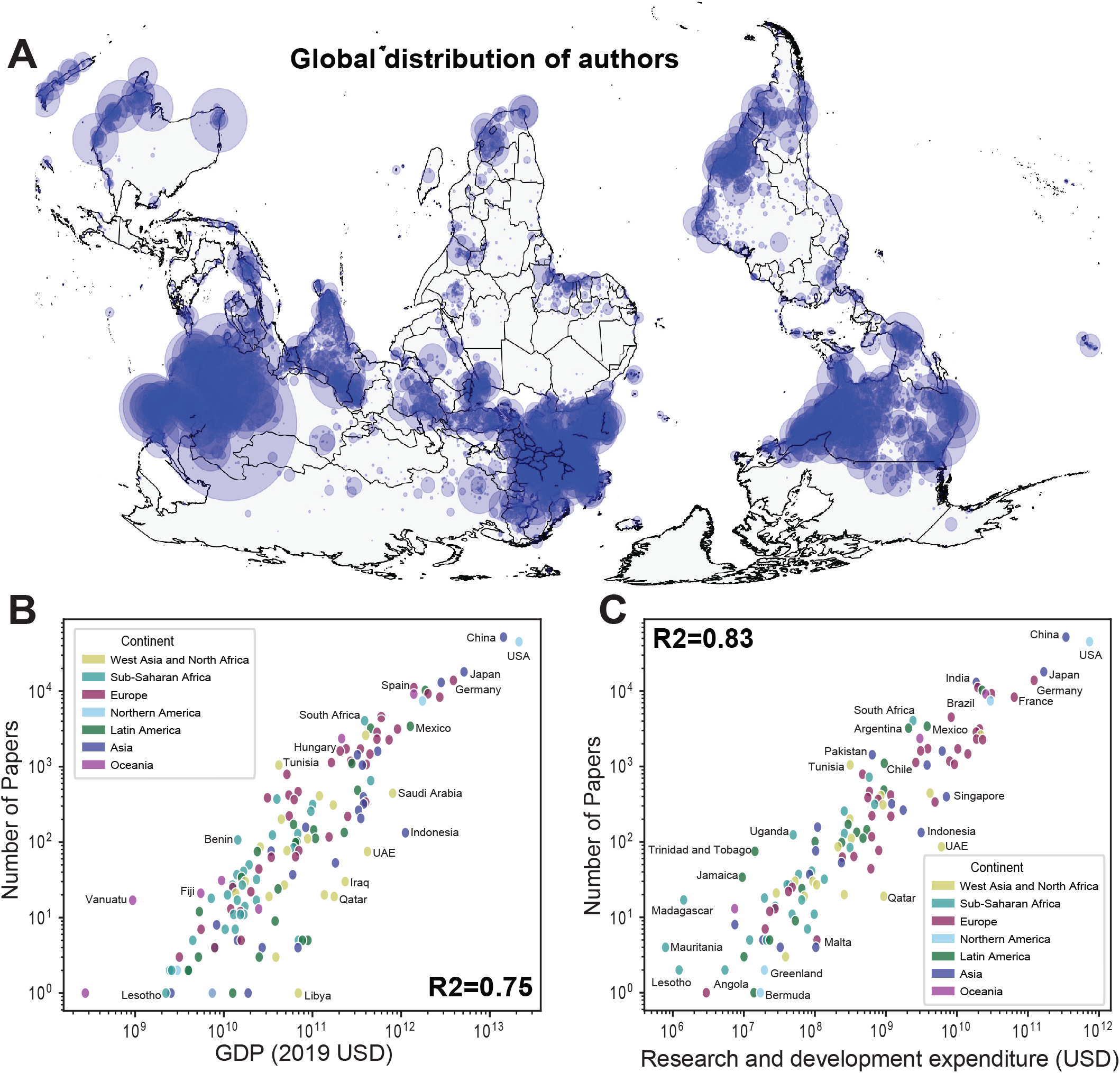
Global patterns of plant science publishing. A) The global distribution of where authors are based, scaled by the number of publications from each location. B) The number of studies published by each nation relative to national Gross Domestic Product (GDP). C) The number of studies published by each nation relative to their research and development expenditure.

However, some individual nations performed better (or worse) than expected. Many emerging economies such as India, South Africa, Mexico, Pakistan, Nigeria, Trinidad and Tobago, Iraq, and Madagascar produced far more publications than expected relative to the money invested in research and development. In contrast, some high income nations, particularly in Scandinavia, Northern Europe, and the Middle East produced far fewer publications than expected relative to the money invested in research and development (Figure 1C and Supplementary Figure 2A). There was very little correlation between publication rate and per capita income (R^2^=0.23, F_2,136_=20, p=1.68e-08) (Supplementary Figure 2B). Based on the United Nations’ income classifications of high, upper-middle, lower-middle, and low income (Supplementary Figure 1B), we found that 61% of all papers published in the last 20 years were led by authors in high income nations. Another 32% were led by authors in upper-middle income nations, and the remaining ~7% of publications were distributed among lower-middle nations. Less than 1% of papers were led by authors in low income nations.

The plant science landscape has changed over the past 20 years. While research output in high income countries has remained relatively stable, there has been a 10-fold increase in the number of papers from upper-middle income nations in the past two decades. In fact, by 2021, there were more papers published by authors in upper-middle income nations than by authors in high income nations (Figure 2A and B). However, this increase was driven primarily by China, which accounted for more than 60% of the publication output from upper-middle income nations in 2020. Other emerging economies such as India, Brazil, Iran, South Africa, Mexico, and Argentina have also made noteworthy contributions to the increased research output of upper-middle income nations (Figure 2C). Publication rates in lower-middle and low income nations have also increased in the past two decades, but still lag far behind those of high and upper-middle income nations (Supplementary Figure 3). In some cases, noticeable decreases in research activity appear to correlate with national disasters and war (e.g. Syria’s annual publications declined sharply in the past 10 years (Supplementary Figure 3A)). Despite noteworthy growth in plant science research, many countries remain underrepresented in the literature.

**Figure 2.**
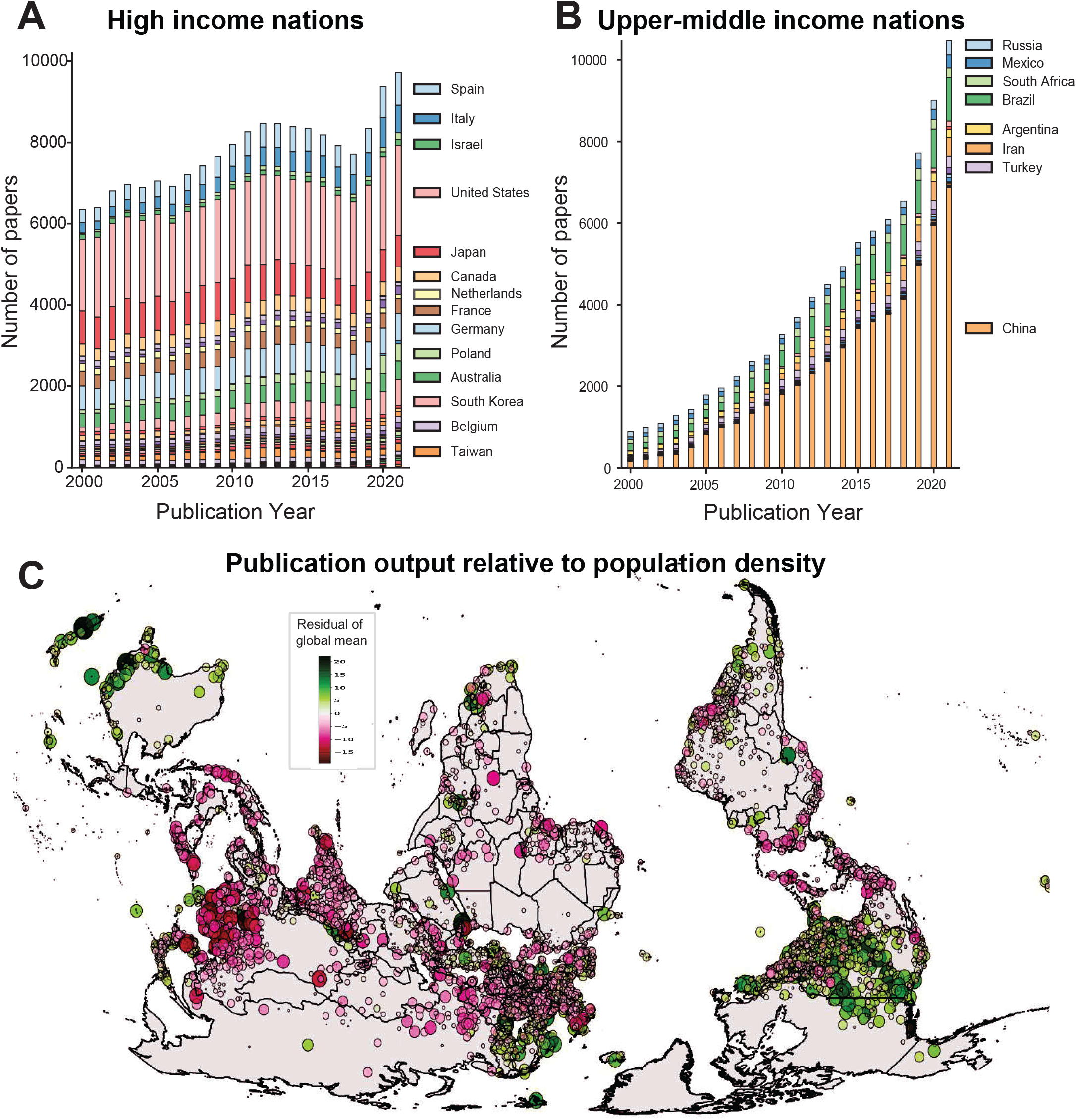
Publication output relative to national affluence and population size. A) The number of studies published each year by authors in high income nations. B) The number of studies published each year by authors in upper-middle income nations. C) Map of publication output relative to population size for locations with more than 300,000 inhabitants or with more than 100 papers produced during the period of 2000-2021. Locations are scaled and colored according to their research output relative to the global trend. Large green points correspond to locations that produce more research than expected based on the global population trend, while large pink circles represent regions that publish less than expected for a city of their size.

In general, productivity is expected to scale with population size following a power law, such that larger cities produce more research output than smaller ones (32–34), and this is what we observed in the plant science literature (Supplementary Figure 4). However, this scaling was variable across the globe. In general, cities in Northern America, Northern Europe, and Oceania had above average research output relative to population size. In contrast, cities in Asia, Africa, and Latin America had below average research output relative to their population size (Figure 2A). Taken together we find that high income nations produced a higher proportion of their research in rural areas, whereas lower income nations concentrated research activity in high-density, urban areas. This is noteworthy because high income nations in Northern America, Europe and Oceania account for less than 10% of the rural population globally (Supplementary Figure 5) but produce more than 64% of the plant science research.

International and intercontinental collaborations were strikingly uncommon in the past two decades of plant science research (Figure 4 and Supplementary Figure 6). More than two thirds (71%) of the publications in our database were written by authors based in a single nation. Just 22% of studies involved a collaboration between two nations, and only 5% of studies included three nations. Less than 1% of studies involved four nations even though 71% of papers have four or more authors, and just 0.04% included five nations despite the fact that 54% of papers had five or more authors. When international collaborations did occur, they tended to be across continents rather than within continents. Only Europe-based authors showed a high frequency of within-continent collaboration (Supplementary Figure 6). Collaborations across continents did occur but were not evenly distributed. Most nations preferred to collaborate with researchers in Europe, Northern America, or China (Figure 3), and were less likely to collaborate with authors in Latin America, Africa, or West Asia. A similar pattern is evident when considering income groupings–only the most affluent nations participated in within-group collaborations, and all other nations preferred to collaborate with high income nations (Supplementary Figure 6).

**Figure 3.**
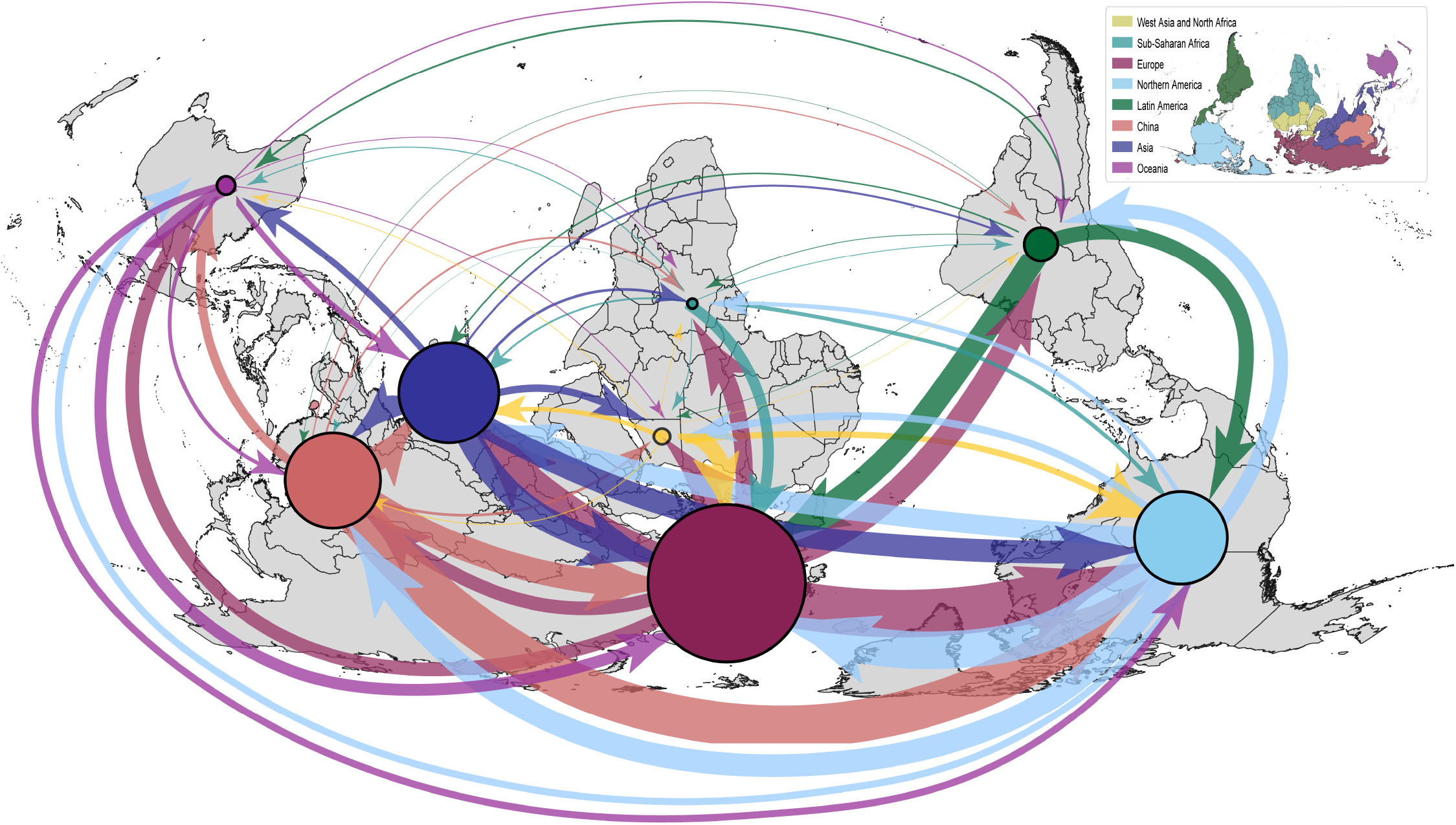
Disparities in global collaborations within plant science research. Circles represent publications that did not involve an intercontinental collaboration. Arrows represent cross-continental collaborations and are directed from corresponding author to co-author. Circles and arrows are scaled by the number of publications.

**Figure 4.**
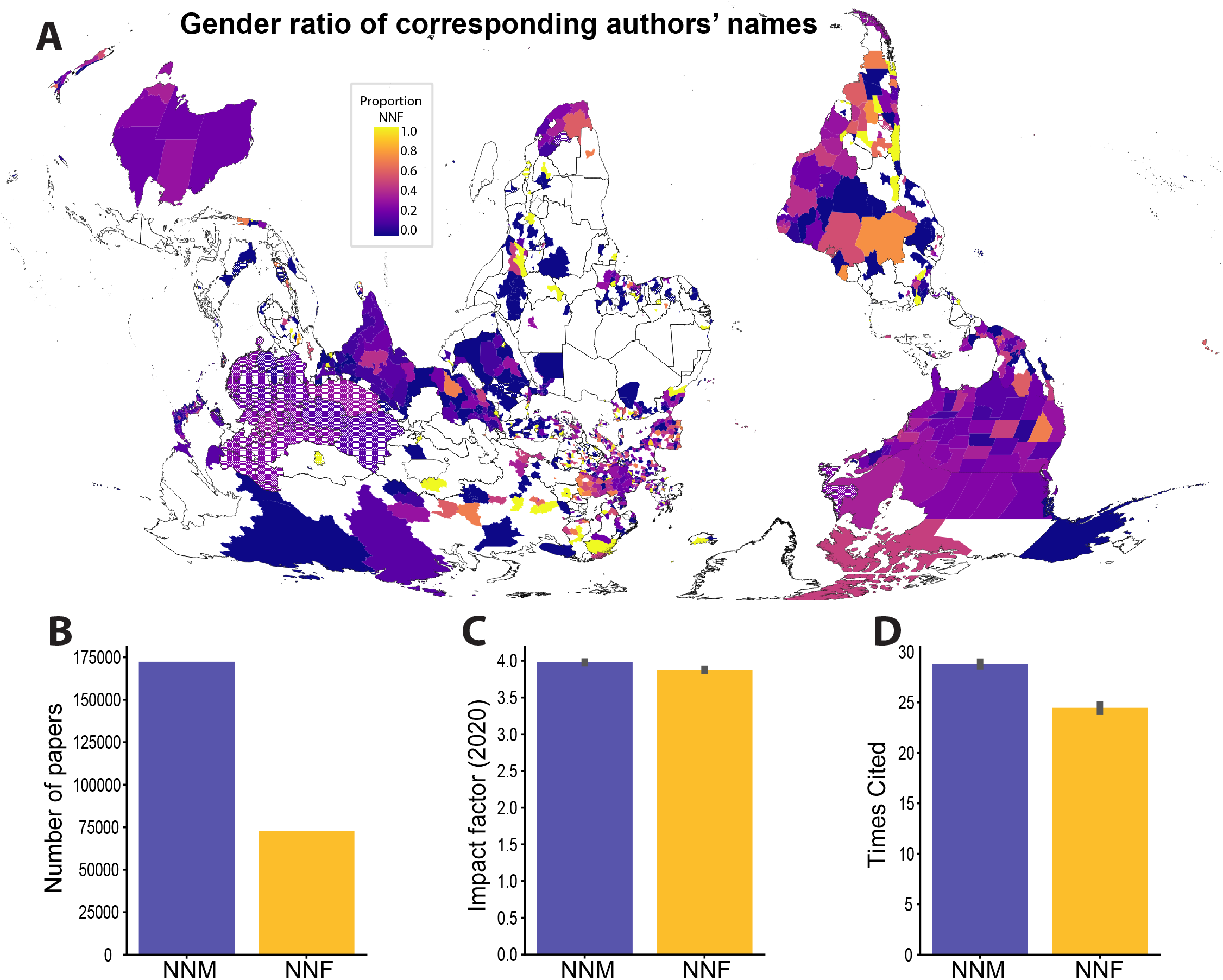
Disparities in the global distribution of corresponding authors by gender. A) Map showing the distribution of name-based gender ratio. All regions where accuracy of name-based gender prediction is less than 90% are faded out. There were many more papers led by authors who had names normatively associated with masculinity (NNMs) than by authors with names normatively associated with femininity (NNFs), but the extent of the imbalance was variable across the globe. B) The total number of publications led by authors with NNMs and NNFs, C) the impact factor of the journals that NNMs and NNFs published in, and D) the citation rates for papers led by authors with NNMs and NNFs.

Despite striking differences in research output, the mean impact factor of the journals that papers were published in spanned just over one point across continents–ranging from 2.92 ± 0.017 for papers led by authors in Sub-Saharan Africa to 4.06 ± 0.011 in Northern America (Table 1). In contrast, citation rates were substantially more variable across continents. In general, papers from the Global South received dramatically fewer citations than those from the Global North, despite publishing in journals with similar impact factors. For example, mean cumulative citations ranged from 17.82 ± 0.304 for papers led by authors working in Sub-Saharan Africa to 36.75 ± 0.298 in Northern America (Table 1)--a twofold difference. This dynamic has remained relatively stable over the past 20 years, with persistent differences in annual citation rates between continents (Supplementary Figure 7). Some individual nations (e.g., China) have seen improvements in citation rates over time, but most have not.

**Table 1.** Continental averages and standard error for the impact factor of journals that authors published in and mean number of citations that papers received.

| *Continent* | *Mean impact factor* | *Mean cumulative citations* |
| --- | --- | --- |
| *Northern America* | $4.06 \pm 0.011$ | $36.75 \pm 0.298$ |
| *Oceania* | $3.67 \pm 0.022$ | $31.99 \pm 0.621$ |
| *Europe* | $3.94 \pm 0.008$ | $31.21 \pm 0.193$ |
| *Asia (minus China and West Asia)* | $3.53 \pm 0.008$ | $26.32 \pm 0.210$ |
| *North Africa and West Asia* | $3.05 \pm 0.017$ | $23.00 \pm 0.409$ |
| *China* | $4.13 \pm 0.009$ | $21.69 \pm 0.159$ |
| *Latin America and the Caribbean* | $3.13 \pm 0.011$ | $18.57 \pm 0.240$ |
| *Sub-Saharan Africa* | $2.92 \pm 0.017$ | $17.82 \pm 0.304$ |

We also investigated how journal policies such as open access fee and society membership related to participation rates for authors with different identities. Of the 296,447 papers examined, only 14% were published Gold open access. Authors in Northern America and Asia published the highest proportion of open access papers (23% and 18% respectively). In contrast, only 10-15% of papers led by authors based in Africa, Latin America, and West Asia were published open access. Of the 16,641 papers published in “elite” journals (with impact factors above seven), 68% were led by authors in high income nations, compared to 61% overall, and another 15% were led by authors based in China. The remaining 17% were distributed across authors in lower income nations. Citation rates were extremely skewed within these journals. For example, papers led by authors in high income nations were cited 82 ± 0.23 times whereas papers from low income nations were cited only 24 ± 0.86 times–a fourfold difference. In general, society journals did not exhibit any more geographic equality in publication and citation rates than the overall trend. Of the 158,711 papers published in society journals 63% were led by authors in high income nations and these received almost double the number of citations (38.9 ± 0.217) compared to papers led by authors in low income nations (20.3 ± 1.767).

### Persistent gender inequalities in the plant sciences

We quantified the effects of patriarchy and gender discrimination in plant science publishing by associating author names with masculinity or femininity. We acknowledge that a binary gender division is an oppressive concept in itself and that true gender is self-identified (35), and we recognize the proximity of our approach to the harmful practice of gender inference. However, we find that we cannot discuss patriarchy and gender discrimination without employing the concept of gender. We purposefully seek to avoid inferring the gender identity of individuals and instead measure the oppressive effects of patriarchy associated with the names themselves. We focus on the normative association of names with masculinity or femininity to measure these effects, and do not presume to know the true gender identity of authors. We further acknowledge that biases in name-based gender inference can arise from the global diversity of cultural naming systems (36). The accuracy of name-based gender prediction varies considerably across ethnicities and is notably poor for East Asian names (37, 38). This is indicative of yet another layer of bias that has resulted in a eurocentric set of tools and analytical frameworks. Improved algorithms that can handle a diversity of naming conventions and accommodate non-binary gender classifications are needed (39). We have tried to adhere to the five principles for ethical gender inference articulated by (38) still, we struggled with the ethics of algorithmic gender inference within our working group and must acknowledge that in conducting such analyses, we too are culpable in propagating the gender binary anew. Given the caveats and obvious shortcomings of name-based gender inference, we urge all readers to interpret these findings with caution.

We hypothesized that individuals with marginalized gender identities (including women, non-binary, gender non-conforming, trans, and people of multiple sexes/genders) would face barriers to participation in plant science and that these would compound with socioeconomic disadvantages and/or historical oppression to further limit participation by intersectional individuals. We cannot test this hypothesis directly without knowing the gender identities of the authors in the paper, so we aimed to instead measure perceptive discrimination based on sexism, that disadvantages individuals with names normatively associated with femininity (40, 41). To test this prediction, names of corresponding authors for each paper were isolated and classified as either 1) names normatively associated with masculinity (NNMs) or 2) names normatively associated with femininity (NNFs) and used as a proxy for gender.

Globally, there were far more papers led by authors with NNMs than authors with NNFs (Figure 4A and B). However, the degree of gender imbalance varied considerably across continents and nations. Among the 20 nations with the highest publication rates, the most NNM biased nations were Japan (14% NNF), India (21% NNF), Netherlands (23% NNF), Switzerland (24% NNF), and Israel (25% NNF). In contrast, the least NNM biased nations were Poland (61% NNF), Argentina (57% NNF), Italy (41% NNF), Brazil (41% NNF), and Spain (38% NNF). On a continental level, Latin America and Europe had the highest proportions of papers led by authors with NNFs whereas Northern America, Asia, and Oceania had the lowest proportion of NNFs. There has been a modest increase in participation by individuals with NNFs over time, but gender ratios remain far from equal across much of the globe (Figure 5).

**Figure 5.**
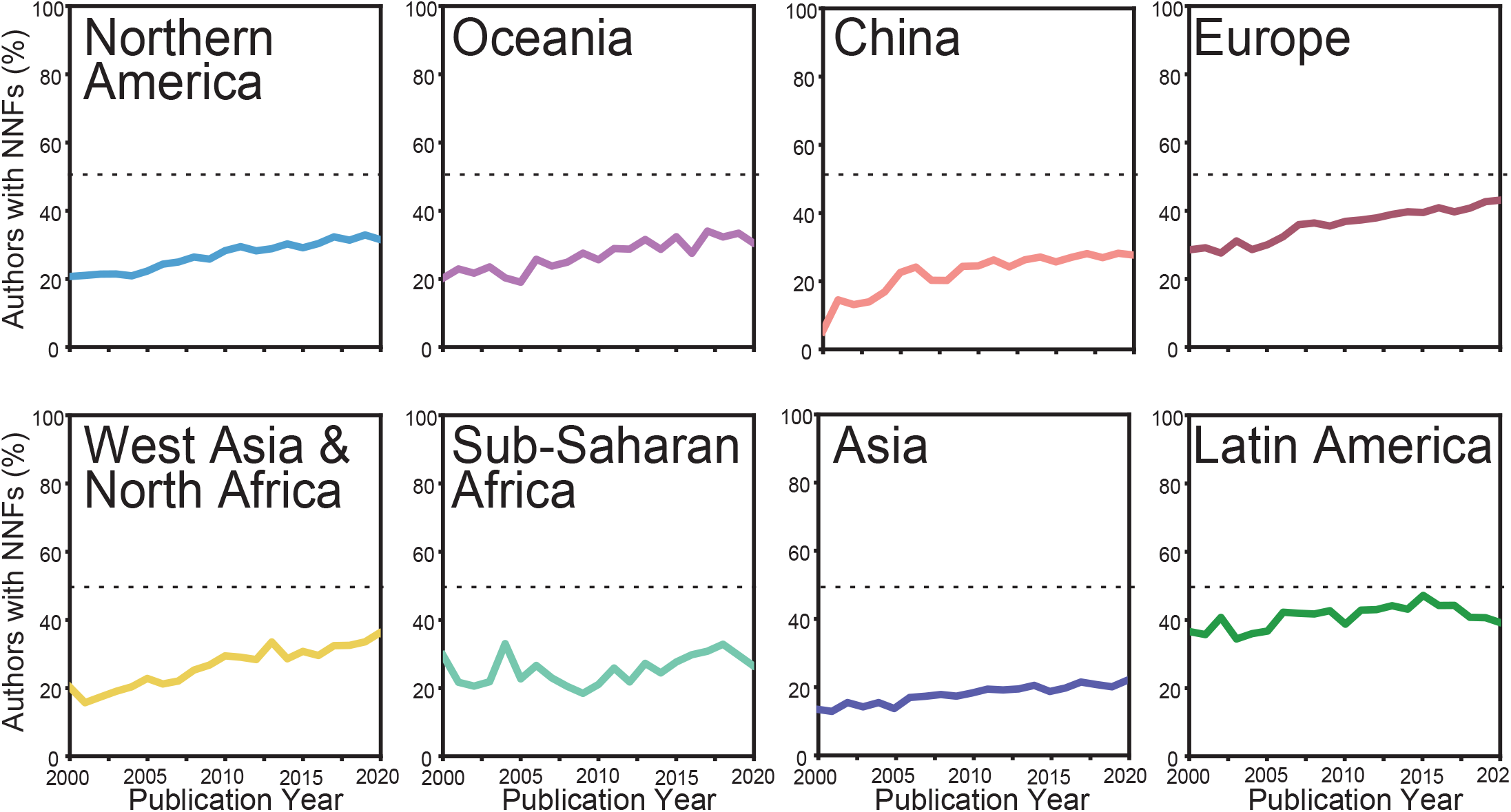
Stagnant gender bias over the past two decades. The proportion of authors with NNFs over the last 20 years is plotted for each of the eight geographical regions investigated.

There was no correlation between national GDP and the proportion of papers led by NNFs (R^2^=0.013, F_2,116_=0.78, p=0.46) (Supplementary Figure 7A). In fact, some of the highest GDP nations had the lowest proportion of NNF authors. There was a similar lack of relationship between per capita income and the proportion of NNF authors (R^2^=0.099, F_2,113_=6.22, p=0.0027) (Supplementary Figure 8B).

There was no significant difference in the impact factor of the journals that authors with NNFs versus NNMs published in. However, there were noteworthy differences in the number of citations these papers received. Papers led by authors with NNMs were cited on average 5 more times than those led by authors with NNFs. This pattern has not improved over time and, if anything, the difference in annual citations for authors with NNFs versus NNMs has expanded (Supplementary Figure 8).

### Taxonomic gaps in focal species studied in the plant sciences

Funding priorities and research activities have historically focused on a narrow subset of described plant species (3, 42, 43) and we expected to find notable taxonomic sampling gaps in the current dataset. To test this prediction, we identified all taxonomic entities mentioned in abstracts via natural language processing. We then summarized overall patterns and geographic differences in the choice of focal species to identify taxonomic sampling gaps and regional patterns.

There were 73,527 unique taxonomic entities represented in our publication database. While the majority of studies focused on plants, we also identified numerous non-plant species including pathogens, symbionts, and other interactors across animalia, fungi, and bacterial groups (Figure 6B). All the top 20 most studied plants represent economically important crop species or models developed by the plant research community (Figure 6A). The model plant *Arabidopsis thaliana* was by far the most studied plant in the past two decades, appearing in four times as many studies as the next most common species wheat (*Triticum aestivum*) (Figure 6A). Poales was the most studied order with over 50,000 mentions, followed by Brasicales, Fabales, and Solanales (Supplementary Figure 10). Many orders were statistically over- or under-represented in the dataset relative to their species richness. The most over-represented orders were Brasicales, Poales, Solanales, Fabales, and Cucurbitales. In contrast, the most under-represented clades were Asterales, Asparagales, Gentianales, Polypodiales, and Lamiales (Figure 6C).

**Figure 6.**
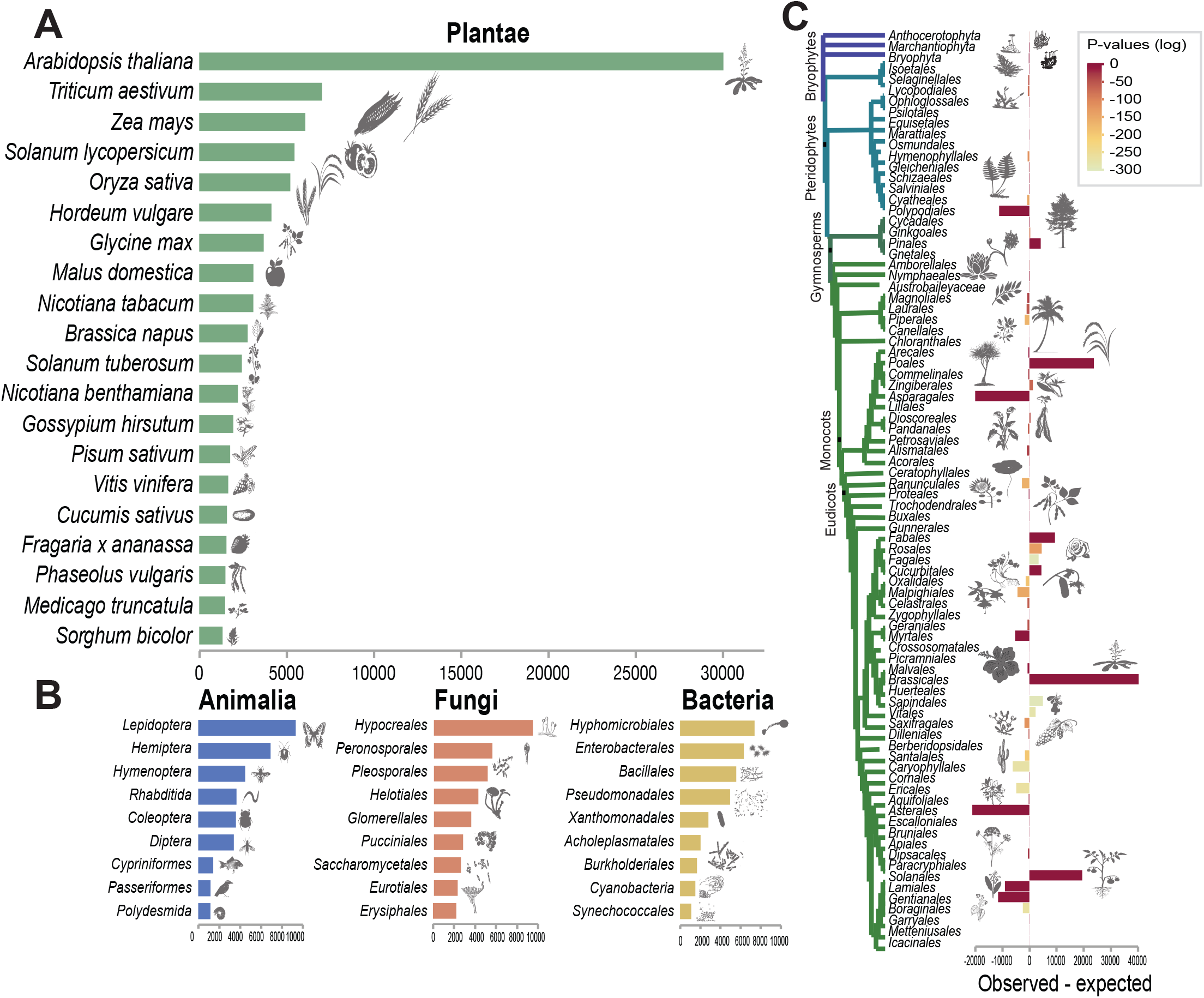
A) The top 20 most studied plant species across all studies. B) The top 9 most studied orders for non-plant groups (animalia, fungi, and bacteria). C) The observed number of studies investigating each order or land plants minus the number expected if sampling effort had been evenly distributed relative to species richness.

We also identified regional differences in the choice of focal organisms. Most high income nations with high publication rates tended to focus on *A. thaliana*, grain crops, vegetables, fruits, and model species (Figure 7). In contrast, many of the nations underrepresented in publishing, tended to focus on lesser-known species and minor or regionally important crops. This finding exemplifies how underrepresentation at the human level impacts the diversity and breadth of focal organisms and research directions.

**Figure 7.**
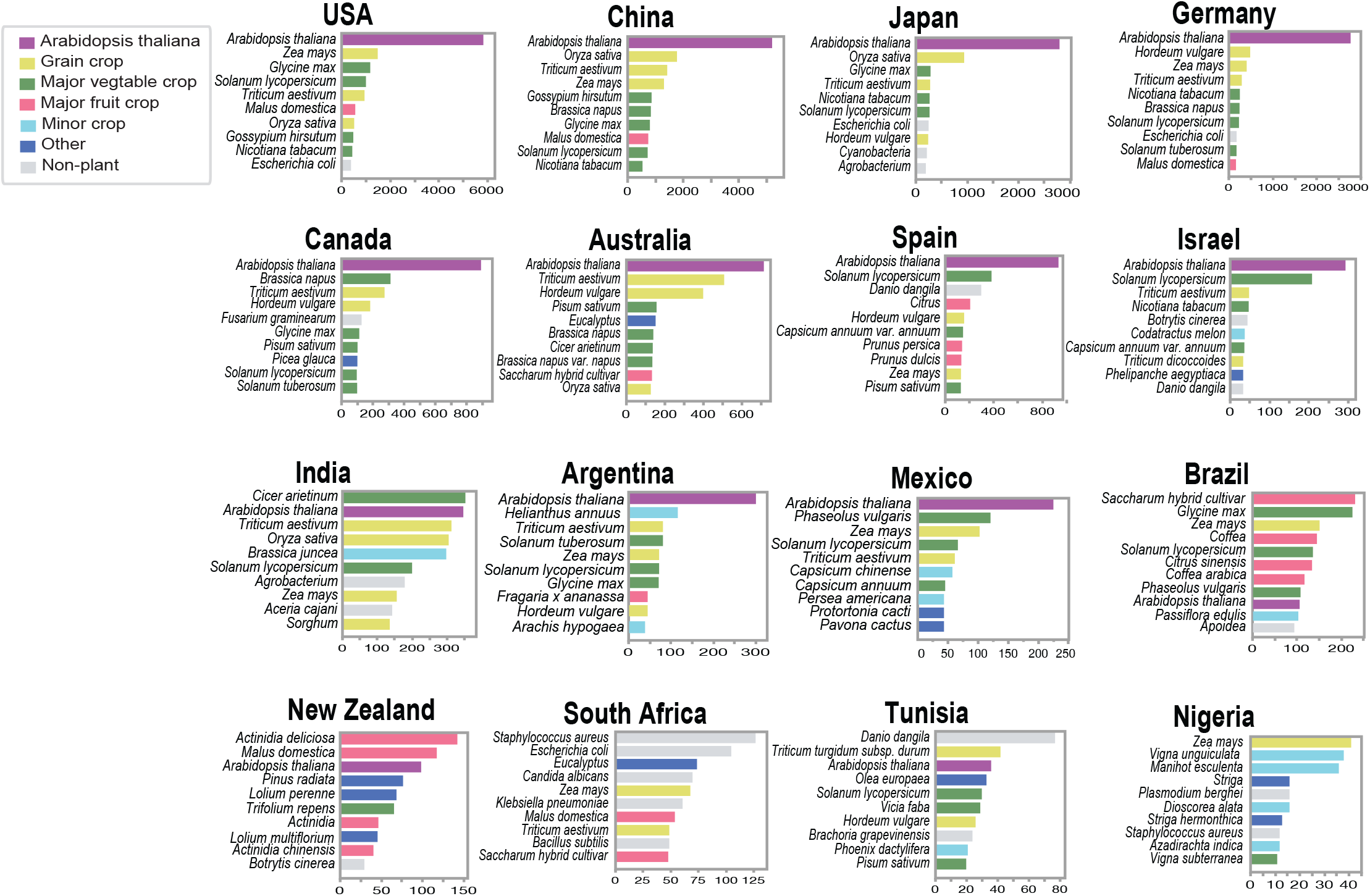
National differences in focal organism choice. The top 10 most studied species from the literature in select nations is plotted. Nations are organized from most prolific to least, and focal organisms are colored by generalized groupings of organism type. The x-axis shows the number of papers that focus on each focal organism.

## DISCUSSION

Our analyses reveal striking geographical biases in plant science research that are associated with national affluence. Global patterns of wealth distribution cannot be understood without an acknowledgement of the impact of imperial colonialism and the resulting consolidation of resources within select nations of the Global North (3, 9, 44). Not only was wealth redistributed during this process, but diverse perspectives and peoples were effectively erased from science as a eurocentric worldview was exported across the globe. Christian missionaries, European traders, and inquisitive researchers all helped to spread frameworks of capitalism, patriarchy, and white supremacy. In biology, these value systems are coupled with a precedence for the English language, reductionist thinking, Latin naming conventions, and biased standards of academic excellence that further exclude individuals from non-European backgrounds. We identified strong correlations between publication rates, GDP, and research and development expenditure. In general, high income nations spent a higher proportion of their GDP on research and development and this led to higher publication output. This finding highlights the privilege of being able to invest in research and development at all, and most lower income nations do not have the necessary funds to support a robust research sector. However, many lower income nations published far more papers than expected relative to their research and development expenditure, while many higher income nations published less than expected. Admittedly, some research output is not captured here because it does not flow into traditional plant science publishing channels and may, instead, be represented by growth in other academic, private, or governmental sectors. Still, this finding gives us reason to pause and recognize the noteworthy accomplishments of scientists from less affluent nations who are doing more with less–an impressive testament to the resourcefulness, creativity, and ingenuity in these regions.

Research output is also associated with increased population density. However, we detected regional differences in this pattern. High income nations (especially in Northern America and Oceania) generated a substantial proportion of their research in rural areas, which makes intuitive sense since plant science is inherently linked to agriculture and natural spaces, and numerous research centers and Land-grant Universities have been built in rural regions. However, lower income nations did not produce many papers in rural areas and instead concentrated research activity in urban centers. We suspect that this pattern is driven by the fact that rural areas are often the last places to be developed and still lack basic infrastructure across much of the globe (45). The differences in rural development impact where research is conducted and contribute to the exclusion of rural peoples and agricultural communities in less affluent nations from the scientific discussion. Only 8% of the world’s rural population lives in Europe, Northern America, and Oceania, but these areas produce more than 64% of the plant science papers. The remaining nations have a disproportionately small publication footprint and the knowledge of ecology, ethnobotany, and agriculture from local and indigenous communities within these areas is largely absent from the literature. These voices and perspectives are often co-opted by researchers from affluent nations through parachute science and other colonialist practices, with no acknowledgment, consultation, or compensation for the discoveries (46). Such gaps in participation undoubtedly translate into gaps in understanding and represent a lost opportunity. These harmful practices have perpetuated persistent inequity in the field.

International and intercontinental collaborations were notably uncommon in the past two decades of plant science research. Of the few international collaborations that we identified, the majority involved a collaborator from Europe, Northern America, and to a lesser degree, China. We suspect that differences in resources (both financial and infrastructural) contribute to these dynamics. Researchers working in high income nations have access to more funding for research, engaging collaborators, and traveling to conferences. Researchers in less affluent nations do not have the same funding opportunities and are therefore limited in the number and type of collaborations they can participate in, as well as the research activities they can undertake. There may also be more subtle and problematic factors driving the skewed collaborative networks we observed. Differences in institutional prestige, born out of eurocentric mindsets, have led some to believe that the best science is done in select institutions in the Global North and that working at or collaborating with those institutions is most desirable. We believe that this rationale is fundamentally flawed and should be dismantled. Affluent nations could do more to engage collaborators in less represented regions of the globe instead of following the well-established global network. Not only would this help to equalize the plant science landscape, but it would enrich our science by bringing in the wisdom of different perspectives.

We identified striking and persistent gender biases in plant science publishing. Given the caveats and shortcomings of name-based gender inference, making specific claims about the gender of individuals or small groups should be avoided, but the overarching patterns identified here are representative. Over 70% of publications in the past two decades were led by authors with masculine names. The extent of gender imbalance was variable across nations and continents but showed remarkably little change over time. In most regions, we detected only modest increases in the number of papers led by authors with feminine names over the past two decades. Interestingly, some of the most affluent nations (e.g., USA, Japan, Netherlands, Switzerland, Germany, Canada, and New Zealand) had extremely male biased publication records despite supposed improvements in women’s rights in many of these nations. In contrast, some less affluent nations in the Global South (e.g., Argentina, Brazil, and Mexico) had among the highest proportions of authors with NNFs. This finding is similar to the “gender-equity paradox” detected in mathematics (47), and contradicts our prediction that individuals facing the intersecting barriers of economic constraints and marginalized gender identity would be more excluded from academic publishing. It suggests that other factors, like cultural differences, could be playing a role in gender inequity. For example, in regions where farming and agriculture are traditionally women’s work, more women may choose to enter the plant sciences. In addition, differences in available support systems can drive career choice, with women sometimes pursuing higher paying jobs (often in STEM fields) when social support systems are limited (48). We looked at a variety of economic development indicators to try to understand what could be driving gender biases in plant science publishing. In contrast to geographical patterns, there was no association between national GDP, research and development expenditure, or per capita income with gender ratio. These findings suggest that the footprint of patriarchy in plant science is deeper than we acknowledge and does not align neatly with narratives about cultural differences in sexism. We also identified gender biases in citation rates that were independent of time, suggesting persistent and ongoing gender discrimination. Because individuals, not institutions drive citation rates, this suggests a deep and pervasive bias running through the discipline. It also means that we, as individuals, have the power to shift these patterns through our actions and choices.

In the past two decades, plant scientists have studied thousands of species spanning plants, animals, bacteria, and fungi. Despite the noteworthy diversity and volume of research, sampling effort has not been equally distributed across clades and taxa. The vast majority of studies have investigated major crop and model species, and the remaining biodiversity accounts for only a fraction of the research on plants. Our analyses identified a number of statistically overrepresented groups of plants, all of which included agriculturally and economically important plants. We also identified numerous underrepresented taxonomic groups, which were ecologically diverse, speciose, and generally of less economic relevance to modern society. These underexplored lineages could provide untold value to humans and ecosystems but have been largely overlooked by modern plant scientists (3, 30, 49). We found some evidence to indicate that taxonomic gaps are related to geographic and gender gaps and we suspect that limited diversity of authors is exacerbating biases in study organism choice. In general, affluent nations in Europe, Northern America, and Asia tended to focus on major crops associated with industrialized agriculture (e.g., wheat, rice, soybean, tobacco, tomato, etc.). In comparison, many of the nations with a smaller footprint in plant science focused their research on regionally important and underutilized crops such as cassava, yam, and millets or local plants with medicinal or historical importance. The disproportionate focus on major crops in the mainstream literature reinforces a homogenization of plant science and limits our ability to conserve and utilize biodiverse plants. It is possible that work on biodiverse species is disproportionately published in regional and subject-specific journals that are not included here, and future studies investigating parallel patterns in these sectors of plant science publishing would be worthwhile extensions of this work. We suspect that if more researchers from across the world were actively engaged in plant science research, there would be a natural diversification of study systems and a broadening of cumulative knowledge.

## Conclusions

Our analyses provide evidence of deep disparities in plant science with links to colonialism, eurocentrism, and patriarchy. Despite the proliferation of statements, committees, workshops, and trainings aimed at increasing diversity, equity, and inclusion, little progress has been made towards actually diversifying plant science in the past two decades (51). These findings can be used as evidence in advocating for change at institutional and policy levels, while also motivating individuals to make positive change in their own research activities and philosophy. While many recognize that the current system is unfair, there are contrasting views on what changes should be made. Some advocate for reformation while others favor abolition, but both agree that there is a need to broaden science and embrace the diversity of knowledge acquisition systems that exist globally. We suggest that first steps towards improving the discipline should consist of a fundamental broadening of our definition of what science is and who can do it. By embracing a more nuanced and context dependent view of data, acknowledging that novelty is not the only source of scientific merit, and recognizing the value of qualitative research, we can begin to minimize colonial biases in academic culture, language, and institutions (30). Funding is another important component, and wealthy nations should take the lead in making efforts to equalize disparities in national affluence established through colonialism. Grants that specifically promote intercontinental collaborations coupled with direct funding to lower income nations could play an important role. Formal policies that provide guidelines and regulations for data ownership and benefit sharing can also help to ensure equitable research practices. The Nagoya protocol represents one such effort, but many nations lack the necessary infrastructure and institutional support to implement the policy effectively. Given the longstanding disparities that exist in plant science, it may be useful to employ concepts of restorative justice, truth and reconciliations practices (50, 51), and a more general shift away from gatekeeping policies and towards inclusive groundskeeping concepts (52). By expanding our definition of what constitutes scientific inquiry and who can take part in it, we begin to open the door to new sources of knowledge. After centuries of centering patriarchal ideals and eurocentric ways of knowing, it is time to make space for other systems of knowledge to rise to the forefront. We hope our analyses can be used to support these positive changes.

## METHODS

### Data acquisition and filtering

We assembled a large-scale database of plant science papers from 127 journals spanning a range of impact factors, nationalities, and sub-specialties (see Supplementary Dataset S1). We cross referenced plant science journals listed in the Journal Citation Reports Database (https://jcr.clarivate.com) with a list of plant science journals compiled by the American Society of Plant Biology (https://plantae.org/plant-biology-journal-database/). We then filtered journals on the following criteria: (1) the journal must have an impact factor, (2) it must be plant specific, and (3) it must include research articles. Metadata associated with all research papers from the resulting 127 journals across the last 20 years were included in the current study. Other metadata were incorporated by referencing JCR and journal webpages, the World Bank 2019 database, the UN Statistics Division, and the UN Department of Economic and Social Affairs (see Supplementary Appendix 1 for methodological details).

### Geography based analyses

The location of authors was inferred from the addresses listed in the papers using an ad-hoc text processing script. Geographic coordinates (geocoordinates) for all these locations were obtained using the Google Maps Geocoding API (https://developers.google.com/maps/documentation/geocoding) with Python via GeoPy. We computed national summary stats, global patterns of author location, and associations with national development indicators using Python (v3.8.8) packages Pandas (v1.5.0) and Numpy (v1.22.4) and visualized data in Seaborn (v0.11.1) and Matplotlib (v3.6.1).

We quantified patterns of collaboration by identifying the location of the corresponding author relative to all other authors for each paper. We then determined if authors were from different countries, continents, or income brackets and summarized global patterns (see Supplementary Appendix 1 for methodological details).

### Gender analyses

We quantified the effects of patriarchy and gender discrimination in plant science by associating author names with masculinity or femininity. The analyses presented here do not identify the true gender of authors. Rather, they show the assumed gender based on the association of first name with either masculinity or femininity. These analyses also likely mis-identify and fail to account for non-binary, gender neutral, and trans individuals, among others. Geographic biases in the performance of gender inference algorithms have also been documented, with most tools performing poorly on East Asian names. This is noteworthy since many of the papers in our dataset are led by individuals with East Asian heritage. Given these caveats, we selected the most robust tool available for this type of analysis (GenderAPI) based on the extensive benchmarking and comparative analyses presented in (36, 37). Summary stats, regional patterns, and changes over time in gender ratios were computed using Python (v3.8.8) packages Pandas (v1.5.0) and Numpy (v1.22.4) and visualized in Seaborn (v0.11.1) and Matplotlib (v3.6.1) (see Supplementary Appendix 1 for methodological details).

### Study Species analyses

The species studied in each paper were identified from abstracts using the Python package TaxoNERD (53). Each biological entity was assigned to a NCBI taxonomy ID and higher-level taxonomic classifications were extracted with ETE Toolkit (54). We summarized the number of mentions for each species, genus, family, and order of land plants to identify sampling gaps in focal organisms and test for statistically over- and under-representation of focal organisms relative to the species richness of the order (see Supplementary Appendix 1 for methodological details).

## DATA AVAILABILITY

Data associated with this study and a description of data acquisition and curation are deposited in Dryad at https://doi.org/10.5061/dryad.pg4f4qrtb.

## AUTHOR CONTRIBUTIONS

RAM, EJA, SP, ARC, CCR, SMT, JMF, DHC and RV conceived of the study. RAM, EJA, SP, ARC, SMT, contributed to data acquisition and curation. RAM, EJA, and SP conducted data analyses. RAM, EJA, SP, ARC, CCR, SMT, JMF, DHC and RV contributed to data interpretation and conceptual framing of the manuscript. RAM, EJA, and SP drew the figures. RAM wrote the manuscript. All authors edited and reviewed the manuscript.

## ACKNOWLEDGEMENTS

This work was supported by NSF Postdoctoral Research Fellowship in Biology (grant no. PRFB-1906094) to RAM and funding from the Michigan State University Department of Biochemistry and Molecular Biology to SP. Other members of our working group were supported by the USDA National Institute of Food and Agriculture, Michigan State University AgBioResearch, the National Research Foundation of South Africa, Universidad Nacional Autónoma de México, and the Water and Life Interface Institute (NSF DBI grant # 2213983).

## Notes

https://doi.org/10.5061/dryad.pg4f4qrtb

### Competing Interest Statement

The authors have declared no competing interest.

### Summary of Updates

Updated analyses and figures

