## Supplementary appendix for "A critical analysis of plant science literature reveals ongoing inequities"

### SUPPLEMENTARY APPENDIX 1

### *A critical analysis of plant science literature reveals ongoing inequities*

*Rose A. Marks, Erik J. Amézquita, Sarah Percival, Alejandra Rougon-Cardoso, Claudia ChibiciRevneanu, Shandry M. Tebele, Jill M. Farrant, Daniel H. Chitwood, and Robert VanBuren*

### SUPPLEMENTARY METHODS

#### *Data acquisition and filtering*

Metadata for all research articles published in a set of plant science journals (Supplementary Dataset S1) during the years 2000-2021 were downloaded from the Web of Science (WoS) using the batch download tool. For each paper, the downloaded metadata included the *Author Full Names, Article Title, Author Keywords, Keywords Plus, Abstract, Addresses--all authors, Address--corresponding author, DOI, Email Addresses, Researcher IDs, ORCIDs, Funding Orgs, Funding Text, Cited Reference Count, Times Cited–All Databases, ISSN, eISSN, Open Access Designations, Quartile, Publication Year, Volume, Issue, IDS Number, UT (Unique WoS ID),* and *Pubmed ID*. The resulting database was filtered to remove duplicate records, papers without a corresponding author, all book chapters, reviews, proceeding papers, and retracted papers. A total of 296,447 records were retained across all 127 journals. The complete dataset, along with a description of data acquisition and curation are deposited in Dryad at https://doi.org/10.5061/dryad.pg4f4qrtb.

Other metadata were incorporated by referencing JCR and journal webpages, including the *Journal Impact factor (2020), Publisher, Publisher City, Publisher Address, Journal location, Open Access options available,* and *Open access fees (USD)*. We consolidated open access designations into two categories for simplicity. Papers were scored as open access only if they were published gold open access. All other open access designations (e.g., green, bronze, etc.) were not considered open access. For more information about open access designations please see (https://clarivate.com/blog/a-researchers-complete-guide-to-open-access-papers/).

Continental divisions and country assignments were based on a sensible combination of subregions as designated by the United Nations Statistics Division (https://unstats.un.org/unsd/methodology/m49/). Income divisions were based on per capita Gross National Income (GNI) as reported by the United Nations Department of Economic and Social Affairs in June 2019 (1). Data on national development indicators (e.g., Gross Domestic Product (GDP), per capita income, and research and development expenditure were taken from the World Bank 2019 database (https://databank.worldbank.org/source/world-developmentindicators and https://data.worldbank.org/indicator/GB.XPD.RSDV.GD.ZS).

#### *Geography based analyses*

The location of every author was extracted from the address as the city, regional administrative division, and country. We kept track of addresses associated with the corresponding authors and addresses associated with the other co-authors, but we were unable to link individual co-authors to specific addresses, compute how many authors were associated with each address on an individual paper, or keep track of authors with multiple affiliations. Thus, we were forced to consider every individual address as one separate author for the global tally. We elaborated a list of unique locations and tallied the number of repetitions for each across the entire database. For every case, especially for countries, we respected the location provided by WoS data. We however recognize that the full addresses listed by WoS might not match perfectly with the addresses listed in the original papers. For example, all institutions based in Hong Kong and Puerto Rico have addresses that list them as part of China and the United States respectively. Similarly, we respected WoS designation for locations under territorial dispute. For example, Sevastopol was listed as part of Ukraine for some addresses, while it was listed as part of Russia in others. We also respected WoS designation of certain territories as individual countries despite their lack of worldwide recognition, such as Palestine, Kosovo, and French Guiana. Addresses corresponding to countries and territories that are no longer recognized were manually examined. All addresses listed as part of Yugoslavia or Serbia and Montenegro were manually assigned to Serbia, as the institutions listed are physically located within modern day Serbia. Similarly, all addresses from the Netherlands Antilles were assigned to Curaçao. Finally, country names were updated to reflect their most recent name, as in Czechia, eSwatini, North Macedonia, and Türkiye.

Geographic coordinates (geocoordinates) for all author locations were obtained using the Google Maps Geocoding API (https://developers.google.com/maps/documentation/geocoding) with Python via GeoPy. Geocoding API failed to provide geocoordinates for 24 locations, which we extracted manually. Locations that were within 25 km from each other were merged and their tallies combined to account for extended metropolitan areas and name changes. For example, the papers associated with Coyoacán, Astana, and Hyderabad, Andhra Pradesh were merged with those from Mexico City, Nur-Sultan, and Hyderabad, Telangana respectively. These merges were performed only when both locations were within the same country to avoid merging cities located at national borders.

We computed national summary stats and associations with development indicators using using Python (v3.8.8) packages Pandas (v1.5.0) and Numpy (v1.22.4) and visualized the data in Seaborn (v0.11.1), Matplotlib (v3.6.1) and Adobe Illustrator (v27.0.1). Briefly, we first computed the number of publications and citation rates for authors working in each nation, continent, and global designation. We then summarized parallel patterns across income levels (as reported by the UN Department of Economic and Social Affairs). Next, we computed both linear and logarithmic regressions to quantify the correlation between publication count and GDP, per capita income, and research and development expenditure (as reported by the World Bank 2019).

To understand how population size relates to publication rate, each location was associated with the population of the closest city listed in the GeoNames database (https://www.geonames.org/). We also added the population of any other city within 25 km to account for a broader metropolitan area. Additional entries and census data were manually added to our local copy of the GeoNames database for locations with no assigned population. The relationship between papers published and population size was computed by first logtransforming the data and then fitting a reduced major axis (RMA) linear regression (2). RMA was chosen to account for the variability of the population size, the x-axis. The true residuals for

RMA fitting (3) were interpreted as the scale-adjusted metropolitan indicators (SAMIs), as proposed by (4), to identify over-performing and under-performing plant science research locations given their population size. To account better for geographical differences, separate scaling models were computed for countries in each continent as well as each income bracket.

The extent and direction of collaborations were estimated by comparing the location of the corresponding author to the location(s) of all other authors on each paper. We considered the primary location of the paper to be the location of the corresponding author and the locations of other authors to be secondary locations. Using this classification scheme we computed the number of papers led by authors based in each continent, income bracket, and nation. We then identified the proportion of those papers that included a collaborator from a different nation, continent, or income bracket. The resulting collaboration network was visualized in Adobe Illustrator (v27.0.1).

#### *Gender analyses*

To identify the first names of authors, we used natural language processing to extract the name of the corresponding author from the complete list of author names. This yielded first names for ~60% of the papers in our dataset. The remaining 40% of entries consisted of abbreviated names in various forms. We manually curated the abbreviated names to fill in gaps and consolidate authors. We used the following rules to obtain full names from abbreviated names and initials: (1) authors with a matching last name and the same first two initials were considered the same author and given the longest form of the first name; (2) authors with a matching last name, first initial, and the same email address were considered the same author and given the longest form of the name; and (3) authors with a matching last name, first initial and based at the same institution were considered the same author and given the longest form of the name, and (4) authors with a matching first initial and the only one of their last name were considered the same author and given the longest form of the name. If the author identity was ambiguous (e.g., two authors with a shared last name and had ambiguous first initial) we left the field blank. Manual curation allowed us to recover the first names for an additional ~25% of the records in our dataset. We submitted these first names along with the nationality of the author to genderAPI to infer the gender of author names.

The resulting gender predictions were used to compute national gender ratios, changes over time, and differences in citation rates for publications by NNFs vs. NNMs. We tested for associations between national development indicators and gender ratios by running linear and logarithmic regressions, computed summary stats, regional patterns, and changes over time using Python (v3.8.8) packages Pandas (v1.5.0) and Numpy (v1.22.4) and visualized data in Seaborn (v0.11.1), Matplotlib (v3.6.1), and Adobe Illustrator (v27.0.1).

#### *Study Species analyses*

The focal organisms in each paper were identified from abstracts using the Python package TaxoNERD. Briefly, TaxoNERD is a deep learning based named entity recognition tool pretrained on biomedical corpus using transfer learning. TaxoNERD extracts biological entities from text and associates each one with its NCBI Taxonomy ID. We then used the ETE Toolkit (Huerta-Cepas, Serra, and Bork, 2016) to extract the higher-level taxonomic classifications for each NCBI taxonomy ID found by TaxoNERD and summarized the number of mentions for each species, genus, family, and order using Python (v3.8.8) packages Pandas (v1.5.0) and Numpy (v1.22.4).

To identify sampling gaps in focal organisms, we compared the observed number of papers focused on each order of land plants to the number that we would expect if research attention had been evenly distributed across focal organisms relative to the species richness of the order. To do so, we computed the total number of species in each order of land plants based as provided in the Leipzig Catalog of Vascular Plants (v1.0.3) (5) and the Missouri Botanical Gardens Index of Bryophytes (http://www.mobot.org/mobot/tropicos/most/bryolist.shtml). We then ran Fisher's exact tests in R (v.4.1.0) to identify clades with a statistical over- or under-representation of papers focused on them. Summary stats, regional patterns, and changes over time were computed using Python (v3.8.8) packages Pandas (v1.5.0) and Numpy (v1.22.4) and visualized in Seaborn (v0.11.1), Matplotlib (v3.6.1), and Adobe Illustrator (v27.0.1).


**A**

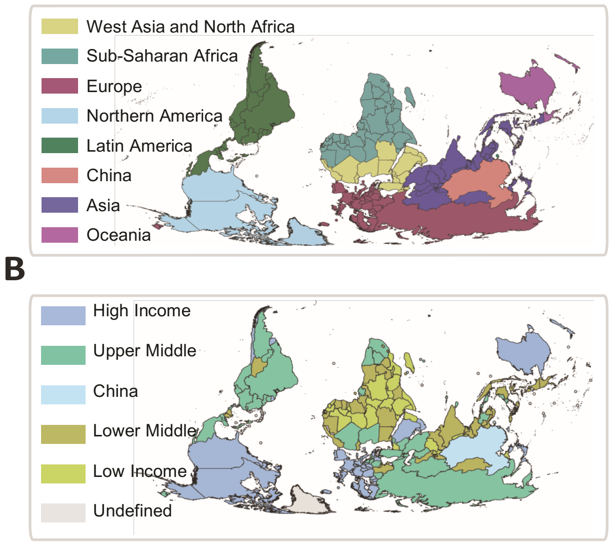


**Supplementary Figure 1. United Nations global designations.** A) Continental regions of the world. B) Income classifications. China is shown separately in both because it is a global anomaly in plant science research.

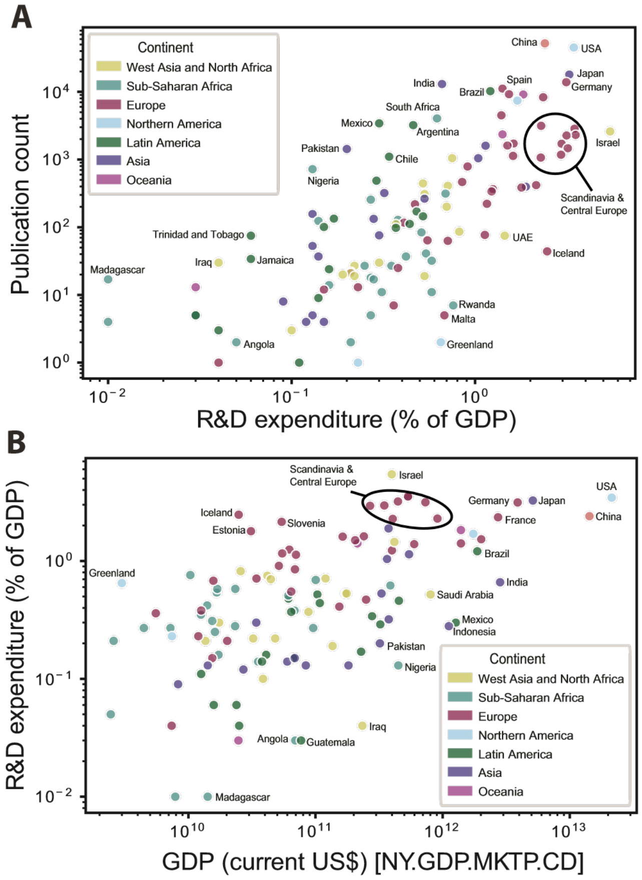


**Supplementary Figure 2.** A) The number of papers published by each nation relative to the proportion of their GDP invested in research and development. B) The proportion of GDP spent on research and development relative to total GDP.

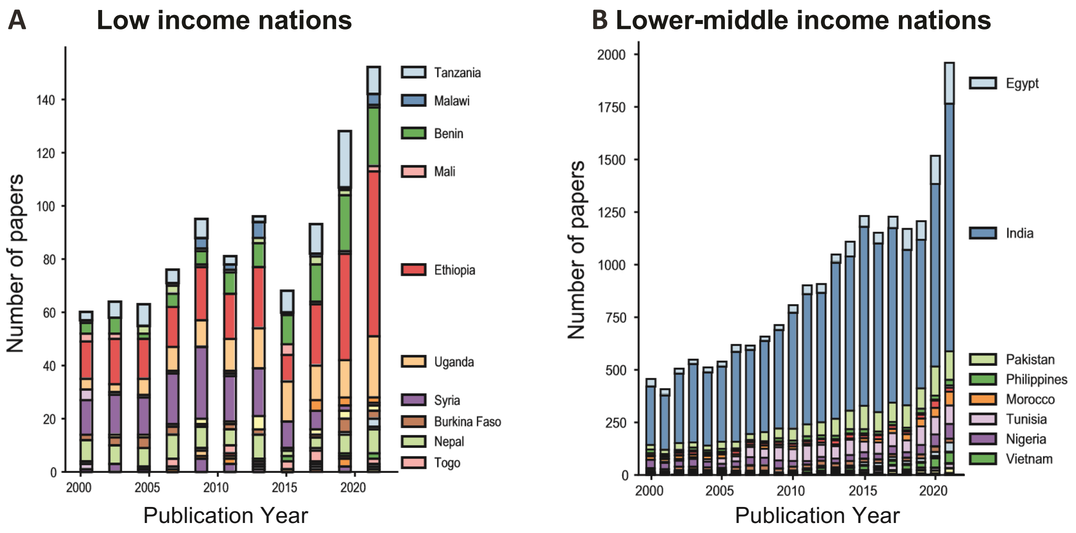


**Supplementary Figure 3. Annual publication rates.** A) The number of papers published each year by authors in low income nations. B) The number of papers published each year by authors in lower-middle income nations.

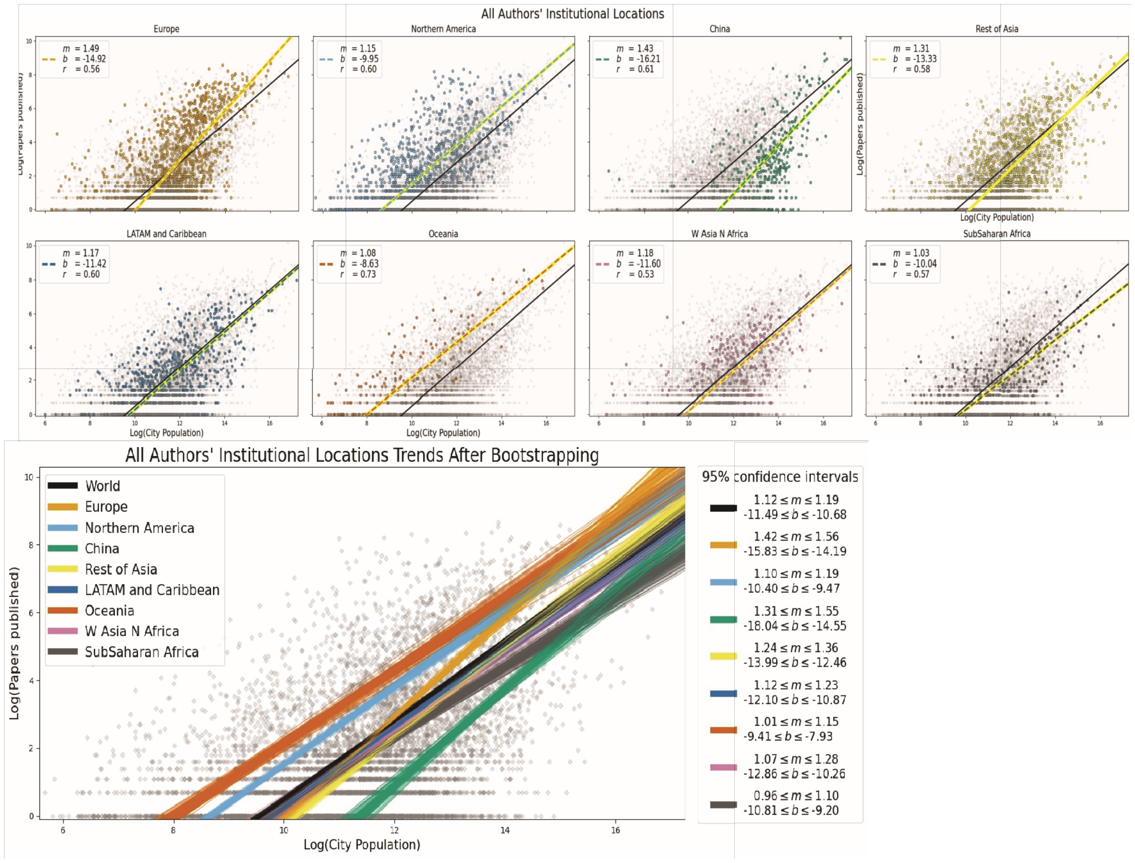


**Supplementary Figure 4. The number of studies published relative to total population by continent**. The black line represents the global trend and the colored lines show continental averages. Bootstrapping plot shows confidence intervals with the spread of the lines representing all the possible trends for that continent.

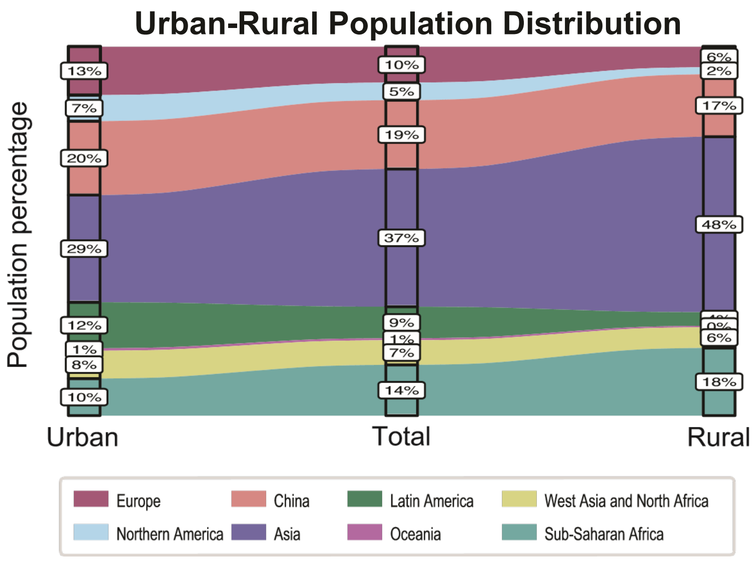


**Supplementary Figure 5. Global distribution of rural and urban populations by continent.** Continental divisions and country assignments were based on a sensible combination of subregions as designated by UN Statistics Division (https://unstats.un.org/unsd/methodology/m49/).

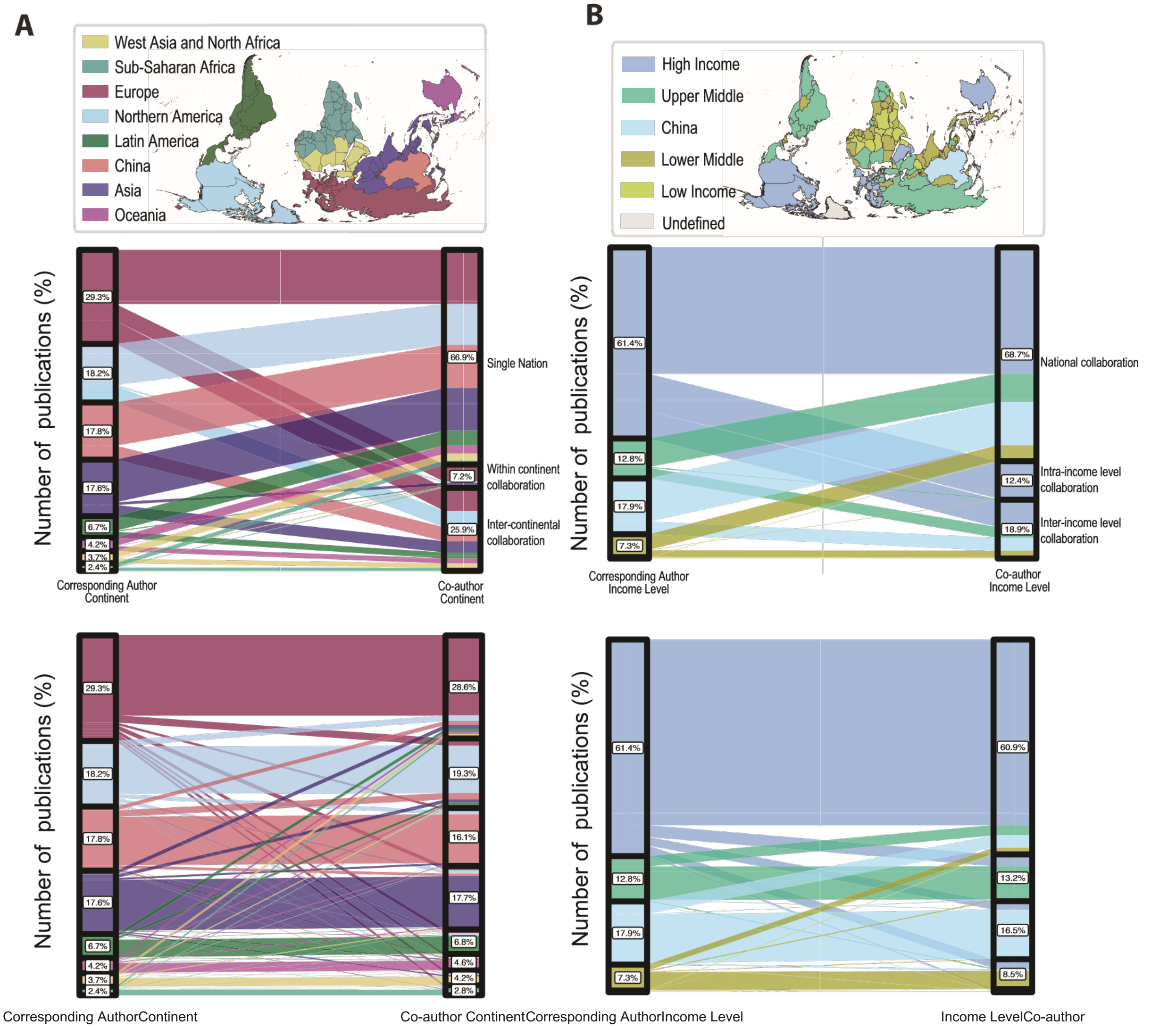


**Supplementary Figure 6. Global collaborations A) across continents and B) income levels.** The continent (or income level) of the corresponding author is shown in the left column and the right column shows the continent (or income level) of coauthors. Flows between columns show the proportion of studies that included authors from each designation. For most studies, all the authors were based in the same nation. When collaborations did span nations there were more likely to be inter-continental than intra-continental. Most continents prefered to collaborate with Europe and Northern America. Similarly, there were few collaborations that spanned income level, and those that did, almost always included a collaborator from a high income nation.

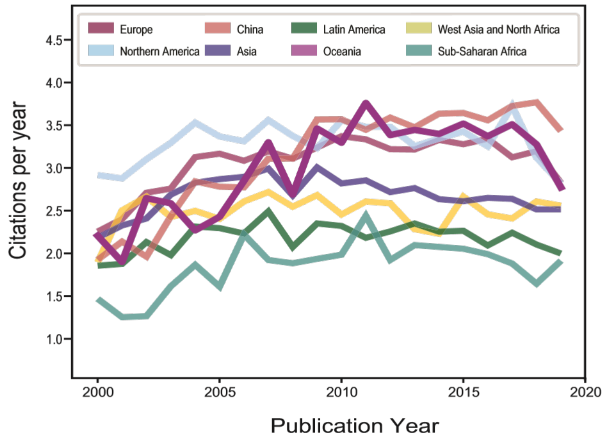


**Supplementary Figure 7. Annual citation rates for eight geographical continents.** China and Europe have experienced improvements in annual citation rates, but most other geographical regions have not.

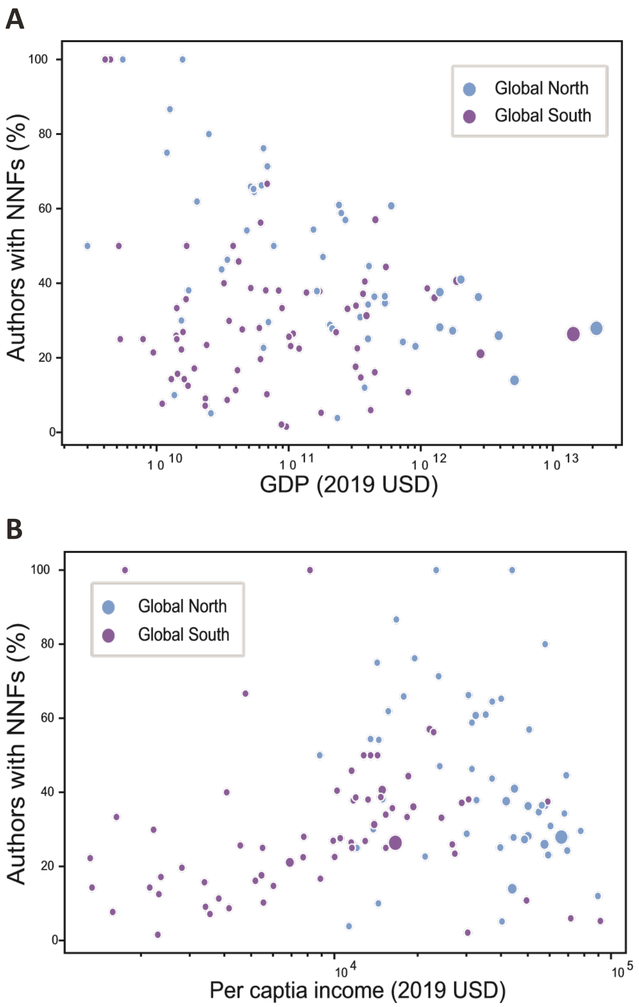


**Supplementary Figure 8.** A) The proportion of papers led by NNFs relative to the gross domestic product (GDP) for all nations. B) The proportion of papers led by NNFs relative to the per capita income for all nations.

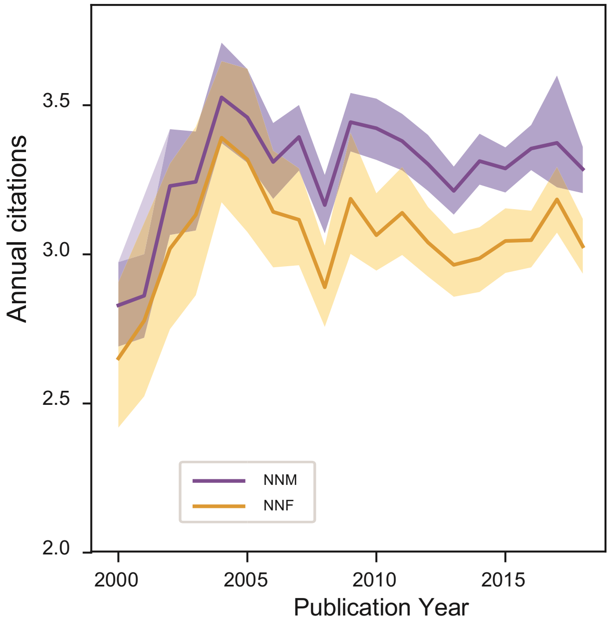


**Supplementary Figure 9.** Annual citation rates for papers led by authors with NNMs and NNFs. Solid lines show the mean and shading represents 95% confidence limits.

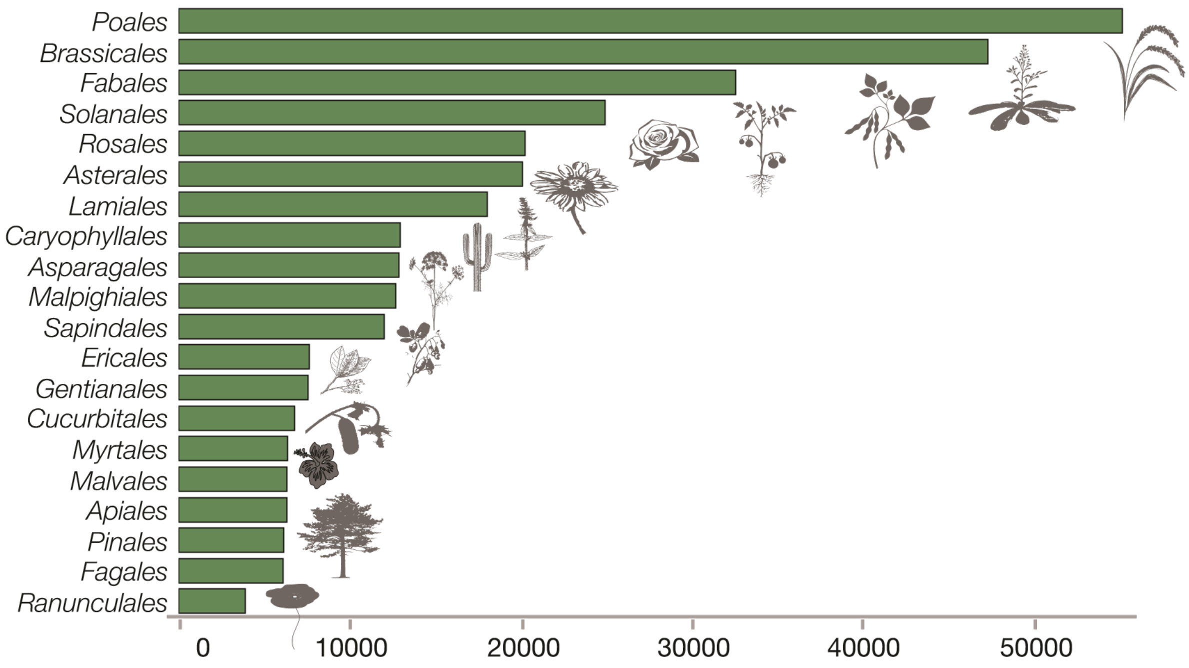


**Supplementary Figure 10.** The top 20 most studied orders of land plants across all studies. The x-axis shows the number of papers focused on that order.
