## Supplementary figures for "A critical analysis of plant science literature reveals ongoing inequities"

### *Global disparities in plant science: a legacy of colonialism, patriarchy, and exclusion*

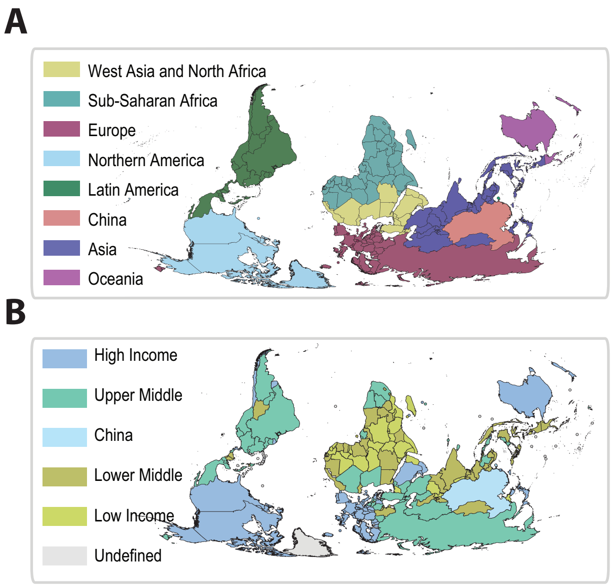


**Supplementary Figure 1. United Nations global designations.** A) Continental regions of the world. B) Income classifications. China is shown separately in both because it is a global anomaly in plant science research.

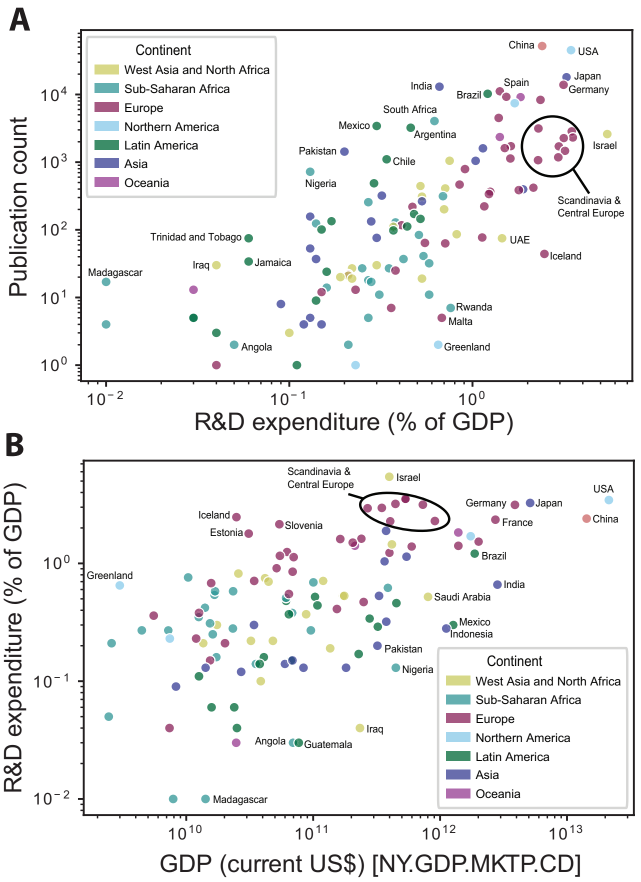


**Supplementary Figure 2.** A) The number of papers published by each nation relative to the proportion of their GDP invested in research and development. B) The proportion of GDP spent on research and development relative to total GDP.

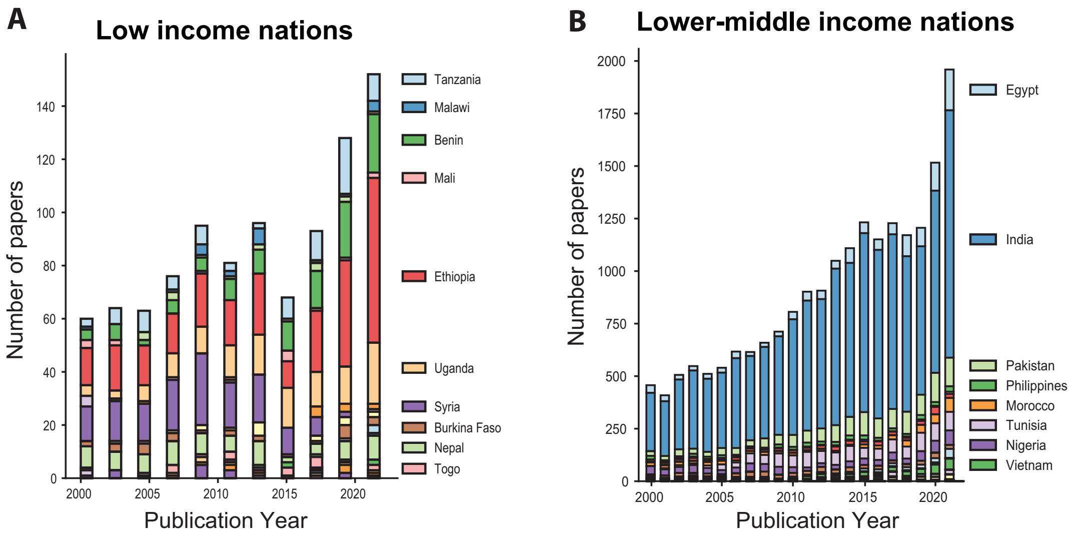


**Supplementary Figure 3. Annual publication rates.** A) The number of papers published each year by authors in low income nations. B) The number of papers published each year by authors in lower-middle income nations.

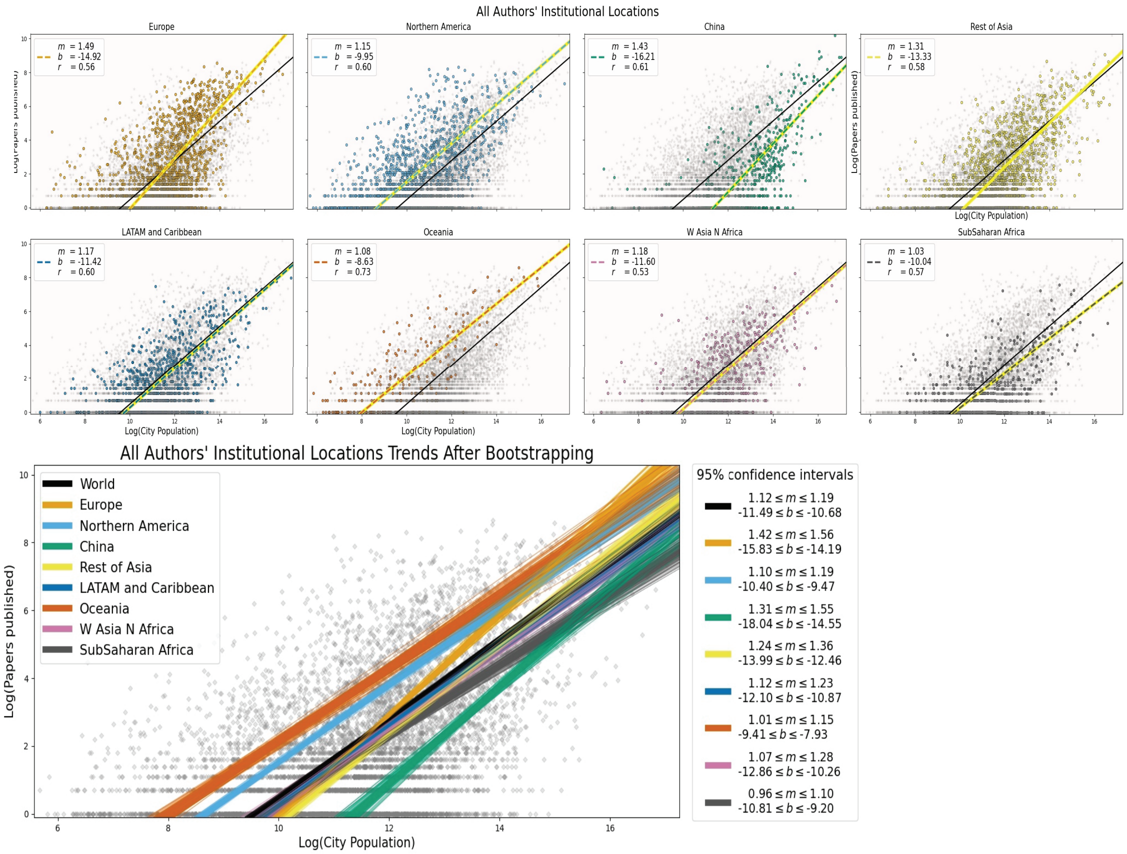


**Supplementary Figure 4. The number of studies published relative to total population by continent**. The black line represents the global trend and the colored lines show continental averages. Bootstrapping plot shows confidence intervals with the spread of the lines representing all the possible trends for that continent.

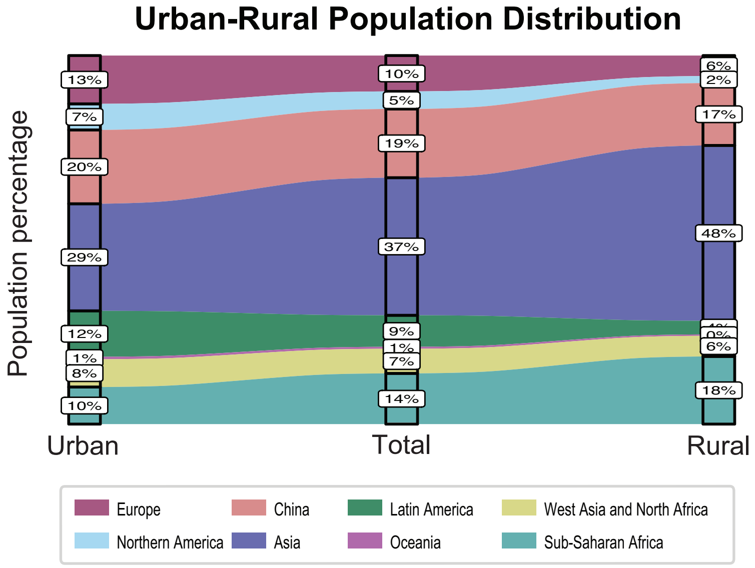


**Supplementary Figure 5. Global distribution of rural and urban populations by continent.** Continental divisions and country assignments were based on a sensible combination of subregions as designated by UN Statistics Division (https://unstats.un.org/unsd/methodology/m49/).

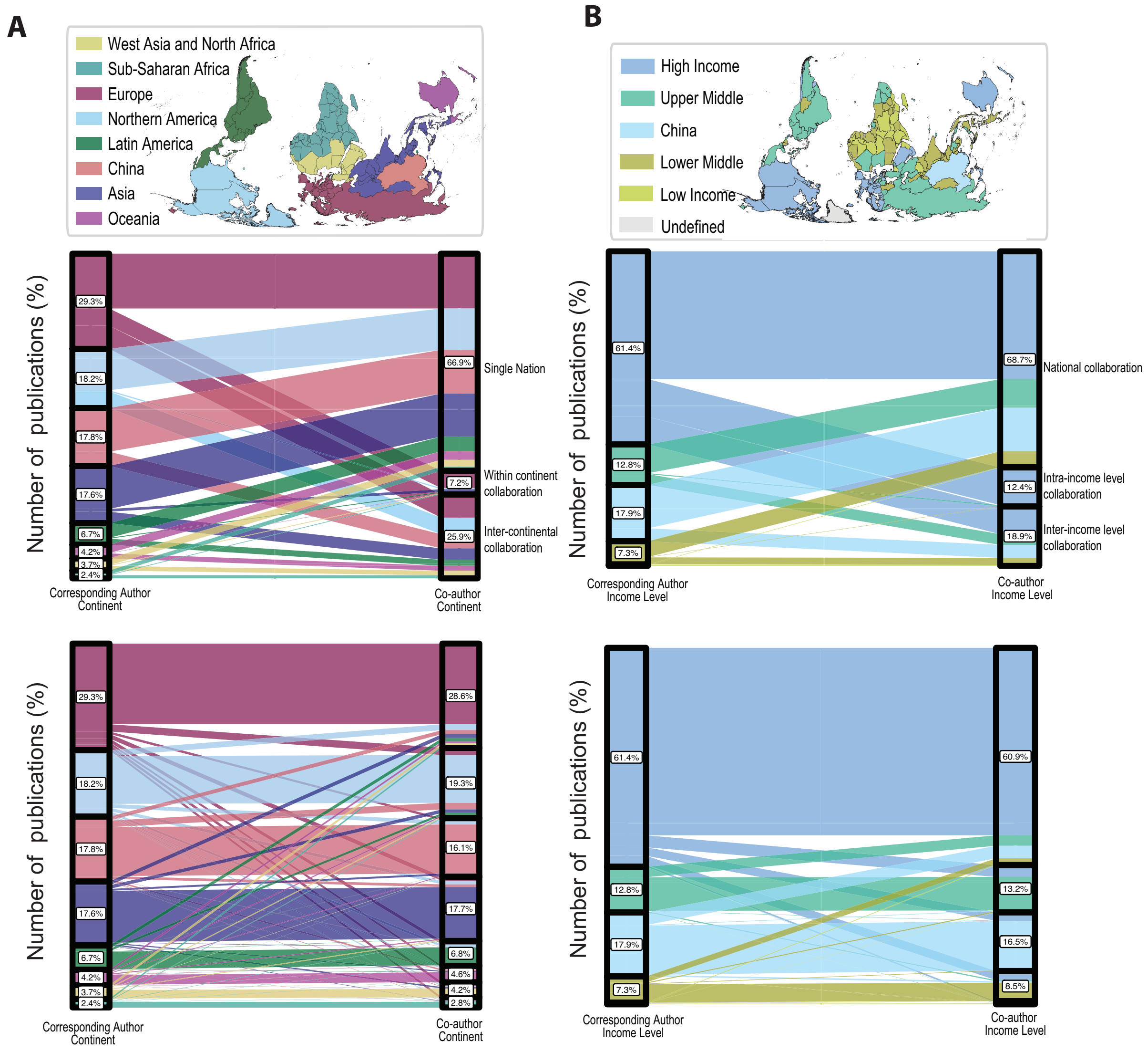


**Supplementary Figure 6. Global collaborations A) across continents and B) income levels.** The continent (or income level) of the corresponding author is shown in the left column and the right column shows the continent (or income level) of coauthors. Flows between columns show the proportion of studies that included authors from each designation. For most studies, all the authors were based in the same nation. When collaborations did span nations there were more likely to be inter-continental than intra-continental. Most continents prefered to collaborate with Europe and Northern America. Similarly, there were few collaborations that spanned income level, and those that did, almost always included a collaborator from a high income nation.

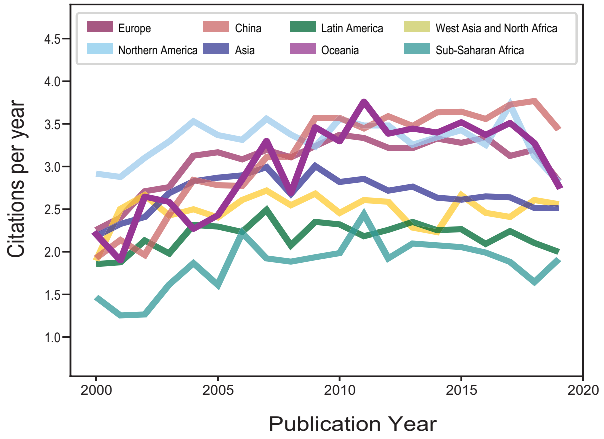


**Supplementary Figure 7. Annual citation rates for eight geographical continents.** China and Europe have experienced improvements in annual citation rates, but most other geographical regions have not.

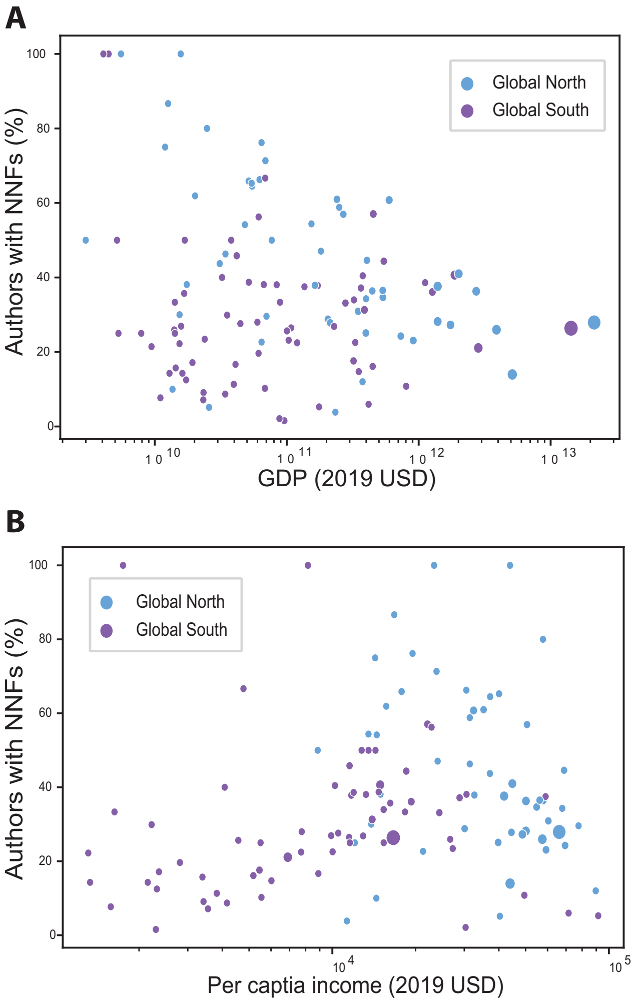


**Supplementary Figure 8.** A) The proportion of papers led by NNFs relative to the gross domestic product (GDP) for all nations. B) The proportion of papers led by NNFs relative to the per capita income for all nations.

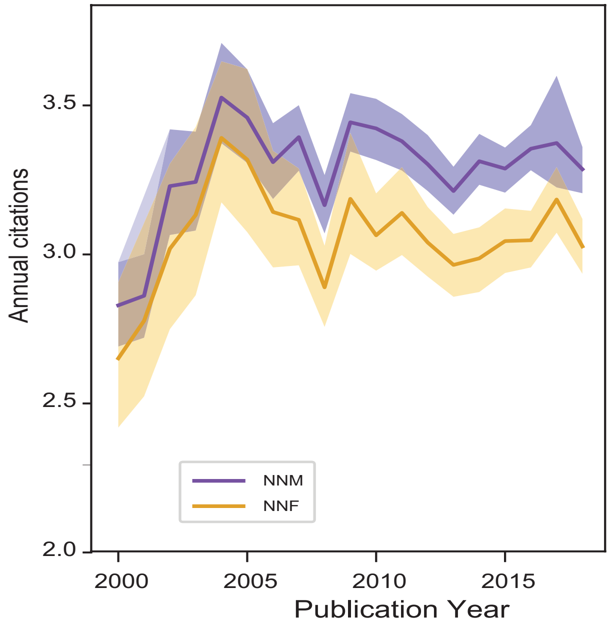


**Supplementary Figure 9.** Annual citation rates for papers led by authors with NNMs and NNFs. Solid lines show the mean and shading represents 95% confidence limits.

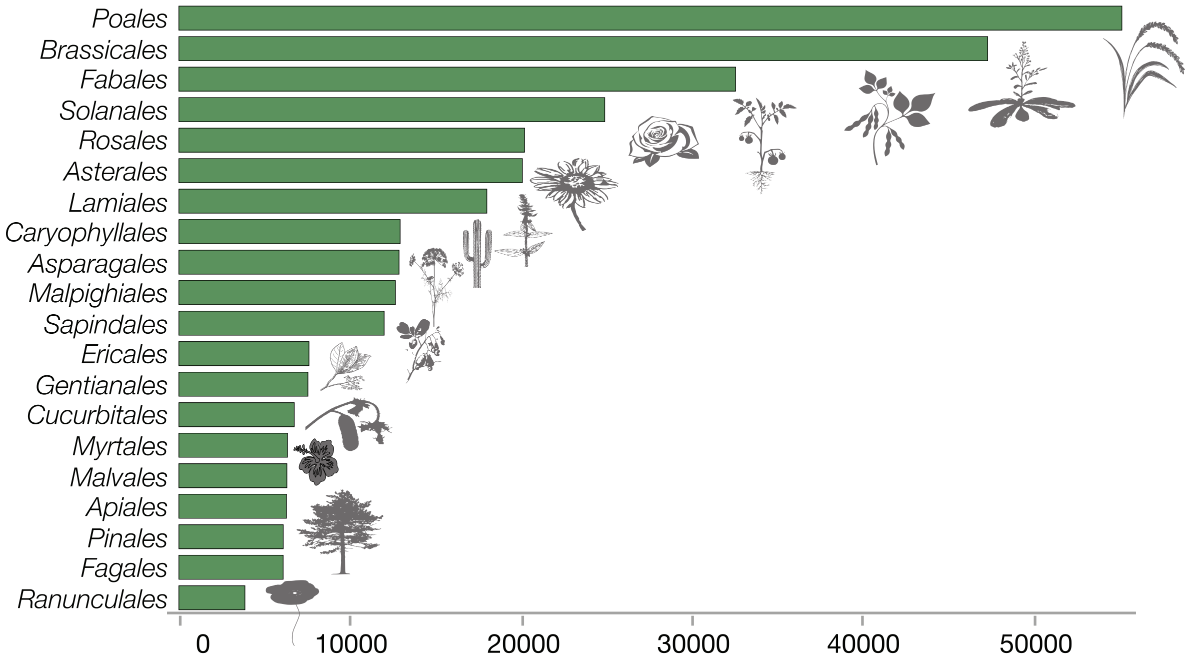


**Supplementary Figure 10.** The top 20 most studied orders of land plants across all studies. The x-axis shows the number of papers focused on that order.
